# PTP1B inhibition promotes microglial phagocytosis in Alzheimer’s disease models by enhancing SYK signaling

**DOI:** 10.1101/2025.05.30.657101

**Authors:** Yuxin Cen, Steven R. Alves, Dongyan Song, Jonathan Preall, Linda Van Aelst, Nicholas K. Tonks

## Abstract

Amyloid-β (Aβ) accumulation is a hallmark of Alzheimer’s disease (AD). Emerging evidence suggests that impaired microglial Aβ phagocytosis is a key feature in AD, highlighting the therapeutic potential of enhancing this innate immune function. Here, we demonstrate that genetic deletion or pharmacological inhibition of protein tyrosine phosphatase 1B (PTP1B) ameliorated memory deficits and reduced Aβ burden in APP/PS1 mice. Moreover, we show that PTP1B was highly expressed in microglia, and its deficiency promoted a transcriptional shift toward immune activation and phagocytosis. Consistently, PTP1B deletion in microglia enhanced phagocytosis and metabolic fitness, supported by increased AKT-mTOR signaling, a pathway essential for meeting the energy demands of activation. Mechanistically, we identified spleen tyrosine kinase (SYK), a key regulator of microglial phagocytosis, as a direct substrate of PTP1B. Inhibition of SYK showed that PTP1B modulates microglial activation in a SYK-dependent manner. These findings established PTP1B as a critical modulator of microglial activation and a potential therapeutic target for AD.

## Introduction

Alzheimer’s disease (AD) is the most common form of dementia and an urgent global issue. As its prevalence rises, the demand for healthcare services grows, placing an increasing burden on society (*1*). AD has a complex pathogenesis, with the Aβ accumulation in the brain recognized as a key initiating event that ultimately contributes to neurodegeneration and cognitive decline, as proposed by the amyloid cascade hypothesis (*2*, *3*). This led to the development of anti-amyloid antibodies, such as aducanumab and lecanemab. Although these treatments help reduce the Aβ burden, their clinical benefits are limited, with concerns regarding potential side effects (*4*, *5*). This suggests a need for novel therapeutic approaches that target multiple aspects of the disease.

In recent years, growing evidence suggests that brain metabolic dysfunction plays a critical role in AD pathology (*6*). Impaired cerebral glucose uptake and utilization may precede cognitive dysfunction by years, or even decades (*7–9*), suggesting this brain hypometabolic state contributes to AD progression. Additionally, systemic metabolic disorders, such as obesity and type 2 diabetes, are well-established risk factors for AD (*10*, *11*). These conditions are associated with chronic inflammation, insulin resistance, and mitochondrial dysfunction, all of which may exacerbate AD pathology (*12*). Together, these findings suggest that targeting metabolic dysregulation and brain energy metabolism could open new therapeutic opportunities for AD (*13*). Consistent with this, repurposing antidiabetic drugs, such as insulin and GLP-1 receptor agonists, has shown promising preclinical and early-phase clinical results for AD (*14–16*).

Protein tyrosine phosphatase 1B (PTP1B) plays a central role in maintaining glucose homeostasis and energy balance (*17–20*). It attenuates insulin signaling and has been implicated in the development of insulin resistance (*17*). Moreover, PTP1B directly regulates leptin signaling in hypothalamic neurons (*18–20*). Inhibition or genetic deletion of PTP1B enhances insulin and leptin sensitivity and improves glucose metabolism in models of obesity and type 2 diabetes (*21–23*). Furthermore, PTP1B inhibition restored impaired insulin signaling and improved the behavioral deficits in a Rett syndrome model (*24*), suggesting its potential to alleviate metabolic dysfunction in neurological diseases. However, PTP1B has long been considered a challenging drug target due to its highly polar and conserved catalytic site (*25*). Notably, recent progress in development of allosteric inhibitors has made it possible to target PTP1B selectively, including in the brain. One such compound, MSI-1436, binds the C-terminal regulatory segment of PTP1B and has showed efficacy in preclinical models (*26*), progressing to clinical trials for obesity and type 2 diabetes, which were discontinued due to commercial considerations rather than scientific limitations (*27*). In this study, we utilized a novel MSI-1436 derivative, DPM-1003, characterized in our lab (*23*, *28*), to explore the therapeutic potential of PTP1B inhibition in an AD animal model.

Beyond its role in metabolic regulation, PTP1B has also been recognized as an important modulator of immune cell signaling. It is well established that PTP1B can dephosphorylate the JAK and Tyk2 kinases and inactivate STAT signaling (*29*). In addition, it has been shown to suppress T cell-mediated antitumor activity through effects on JAK/STAT5 signaling (*30*). In macrophages, PTP1B regulates myeloid cell differentiation and activation through CSF1 and IFN-γ signaling (*31*, *32*). Considering that microglia are brain-resident macrophages, these findings prompt a broader immune-regulatory role for PTP1B that may extend to microglia. Interestingly, recent findings suggest that PTP1B is a positive regulator of microglia-mediated neuroinflammation (*33*); however, these findings were observed outside of the context of AD, highlighting the need for further investigation.

Microglia play a key role in AD by mediating chemotaxis and phagocytosis to clear toxic aggregates. Notably, anti-amyloid immunotherapies often rely on microglial activation to clear Aβ plaques (*34*, *35*), highlighting the potential importance of targeting microglial function in AD therapies. Interestingly, microglial activation has been associated with enhanced cerebral glucose uptake in AD mouse models and patients (*34*, *35*), suggesting that metabolic regulation is closely linked to microglial function. Indeed, modulating microglial metabolism directly impacts their functions, such as phagocytosis(*36–40*), suggesting that targeting metabolic pathways could enhance Aβ clearance and mitigate disease progression. Among the signaling pathways, SYK plays a central role downstream of multiple immunoreceptors in microglia, coordinating both metabolic fitness and phagocytosis (*41–43*). Interestingly, a previous study suggested that loss of PTP1B enhanced B cell activation signaling, including the phosphorylation of SYK (*44*). Based on this, we hypothesized a possible connection between PTP1B and SYK signaling in microglia. Considering the role of PTP1B in both metabolic and immune signaling, it has the potential to regulate microglial function at the intersection of these pathways.

In this study, we investigated whether PTP1B could serve as a therapeutic target for AD. Using the APP/PS1 mouse model of AD, we demonstrated that deletion or pharmacological inhibition of PTP1B significantly improved learning and memory and reduced Aβ burden in the brain. In addition, PTP1B deletion triggered a more reactive and phagocytic transcriptional profile in microglia, enhancing their phagocytic activity in vitro and in vivo. Mechanistically, we demonstrated that SYK is a direct substrate of PTP1B, and that this interaction is critical for regulating microglial activation in response to Aβ stimulation. Together, these findings highlight a critical role of the phosphatase in microglial function and suggest that targeting PTP1B may represent a promising strategy to enhance microglial-mediated Aβ clearance and mitigate AD progression.

## Results

### Deletion and inhibition of PTP1B in APP/PS1 mice resulted in beneficial cognitive effects

To test whether PTP1B can serve as a potential therapeutic target for AD, we cross-bred PTP1B knockout mice with APP/PS1 mice (APP/PS1;PTP1B^-/-^). We chose APP/PS1 mice as our AD mouse model for two reasons. First, this model is well-established to study amyloid pathology, as it exhibits age-dependent Aβ accumulation starting at 6-months of age and progressing to cognitive deficits around 12-months (*45*, *46*). Second, this mouse model demonstrates impaired glucose and insulin tolerance as early as 6-months (*47*, *48*).

By 12-13 months of age, these mice were tested for two different behavior tests (fig. S1A). Recognition memory was assessed by the Novel Object Recognition behavior test, which leverages the natural tendency of rodents to explore novel objects over familiar ones (Fig. 1A). Interestingly, PTP1B deletion improved recognition memory in APP/PS1 mice, as shown by increased exploration of the novel object compared to APP/PS1 controls (Fig. 1B). Furthermore, spatial learning and memory were evaluated using the Morris Water Maze (Fig. 1C). During the acquisition trials, PTP1B deletion in APP/PS1 mice shortened the time taken to find the platform (escape latency) compared to APP/PS1 mice (Fig. 1D), suggesting improved spatial learning. Twenty-four hours after the last training session, with the platform removed, PTP1B deletion in APP/PS1 mice increased the number of crossings over the previous location (Fig. 1E) and the time spent in the target quadrant (Fig. 1F), indicating improved spatial memory. No differences in swimming speed were observed among the groups (fig. S1B), confirming that the observed effects were not influenced by variations in motor function.

**Fig 1.**
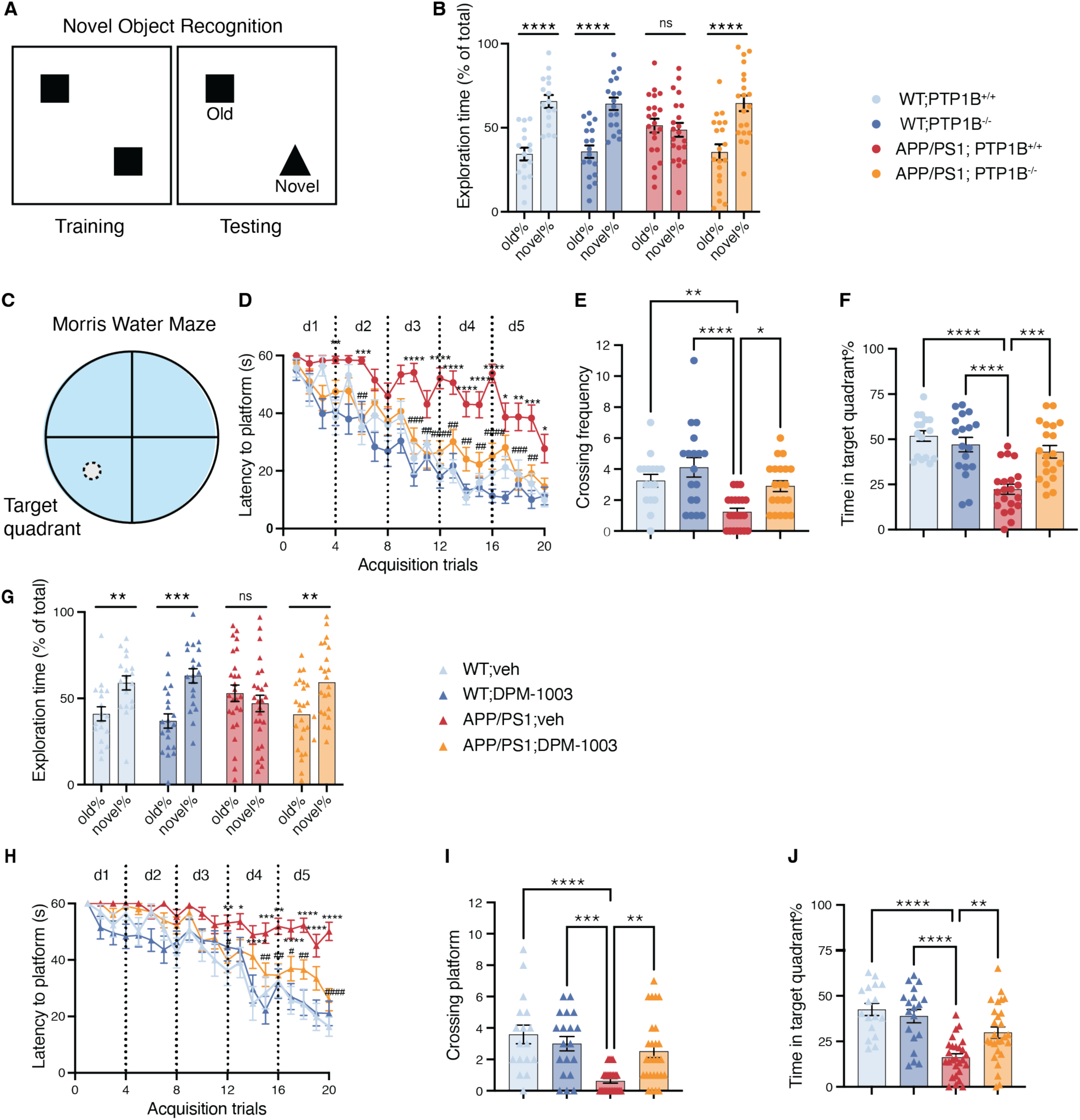
Deletion or inhibition of PTP1B in APP/PS1 model improved learning and memory. (**A**) Schematic of the Novel object recognition tests. (**B**) Percentage of exploration of old or novel object in WT or APP/PS1 mice with or without PTP1B deletion. (**C**) Schematic of the Morris water maze tests. (**D-F**) Morris water maze tests on WT or APP/PS1 mice, with or without PTP1B deletion; (D) Escape latency to submerged platform in acquisition trials (*: comparisons between WT;PTP1B^+/+^ and APP/PS1; PTP1B^+/+^, #: comparisons between APP/PS1; PTP1B^+/+^ and APP/PS1; PTP1B^-/-^); (E) Crossing frequency in the platform area in probe trial; (F) Time in target quadrant in probe trial; the symbol legend in Fig. 1B also applies to Fig. 1D-F. (**G**) Percentage of exploration on old or novel object on WT or APP/PS1 mice treated with vehicle or PTP1B inhibitor DPM-1003. (**H-J**) Morris water maze tests on WT or APP/PS1 treated with vehicle or DPM-1003; (H) Escape latency to submerged platform in acquisition trials (*: comparisons between vehicle-treated WT and mice; #: comparisons between APP/PS1 mice with vehicle or DPM-1003 treatment); (I) Crossing frequence in the platform area in probe trial; (J) Time in target quadrant in probe; the symbol legend in Fig. 1G also applies to Fig. 1H-J.

In addition, we examined the effects of an allosteric PTP1B inhibitor, DPM-1003, in APP/PS1 mice. Starting from 11 months of age, a stage at which the mice exhibit significant amyloid plaque deposition and cognitive deficits, the mice received the compound for 5 weeks (fig. S2A). Similar to gene ablation, PTP1B inhibitor treatment significantly improved the recognition memory, as illustrated by an increased novel object exploration percentage in DPM-1003-treated APP/PS1 mice compared to the saline-treated controls (Fig. 1G). In the Morris Water Maze, DPM-1003-treated APP/PS1 mice showed shorter escape latency (Fig. 1H) and enhanced spatial memory, indicated by increased crossings of the former position of the platform and longer time spent in the target quadrant (Fig. 1I-J). Swimming speed was comparable across all groups (fig. S2B). These results demonstrate that both PTP1B deletion and inhibition alleviate cognitive impairments in APP/PS1 mice.

### Deletion and inhibition of PTP1B reduced Aβ levels in APP/PS1 mice

Since Aβ accumulation is associated with cognitive deficits in APP/PS1 mouse model, we investigated whether PTP1B deletion or inhibition could impact Aβ burden in this model. Thioflavin S (ThioS) selectively labels fibrillar Aβ, whereas 6E10 detects both the fibrillar and non-fibrillar forms (*49*); this dual labeling enables a comprehensive assessment of Aβ burden. Notably, WT animals did not show any plaques (fig. S3A), whereas APP/PS1 mice displayed plaque formation (Fig. 2A). Quantitative analysis in the hippocampus of APP/PS1 mice indicated that PTP1B deletion reduced amyloid deposition. Specifically, the area covered by ThioS-positive plaques was reduced by 33% in APP/PS1 lacking PTP1B mice compared to APP/PS1 controls (1.7 ± 0.11 vs 1.1 ±0.08) (Fig. 2B), and 6E10-positive area was reduced 27% (1.8 ± 0.1 vs 1.3 ± 0.08) (Fig. 2C). Similarly, treatment with DPM-1003 in APP/PS1 mice also led to reductions in Aβ burden, with ThioS-positive areas reduced by 28% (1.5 ± 0.17 vs 1.1 ±0.08) and 6E10-positive area decreased by 26% (1.5 ± 0.12 vs 1.1 ±0.08) (Fig. 2D-F).

**Fig. 2.**
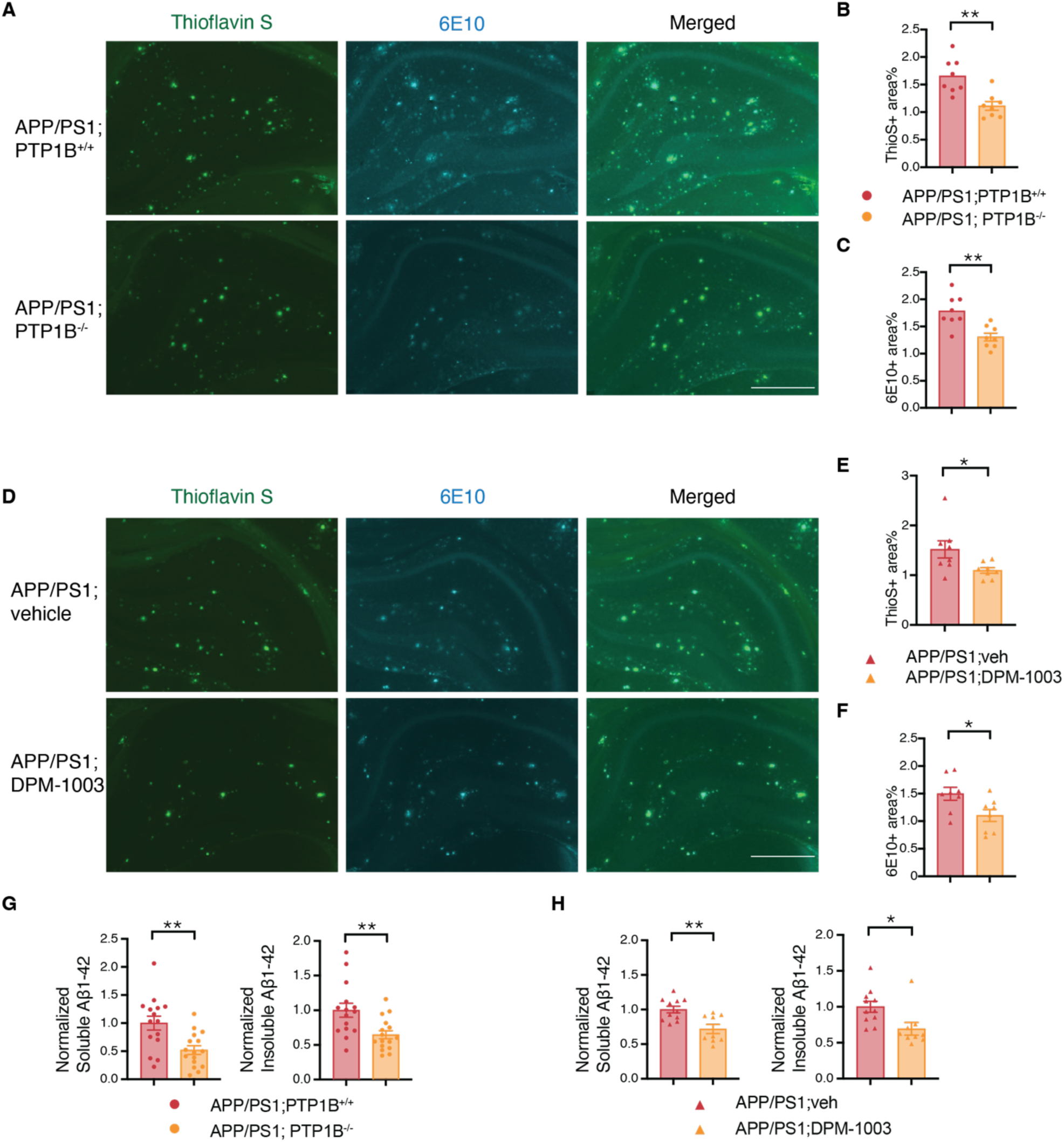
Deletion or inhibition of PTP1B in APP/PS1 reduced Aβ levels in APP/PS1 mice. (**A-C**) Immunofluorescence and quantification of Aβ levels in APP/PS1 mice with or without PTP1B in hippocampal region; (A) Representative images showing Thioflavin S and 6E10 staining, Scale bar =500µm; (B) Quantification of percent area occupied by ThioS; (C) Quantification of percent area occupied by 6E10; n=8/group, 3-4 brain slices/mouse. (**D-F**) Immunofluorescence and quantification of Aβ levels in APP/PS1 mice with or without DPM-1003 treatment in hippocampal region; (D) Representative images showing Thioflavin S and 6E10 staining, Scale bar =500µm; (E) Quantification of percent area occupied by ThioS; (F) Quantification of percent area occupied by 6E10; n=8/group, 3-4 brain slices /mouse. (**G**) ELISA quantification of Aβ_1–42_ levels in the brain diethylamine (soluble) and formic acid (insoluble) fractions of cortical tissue in APP/PS1 mice with or without PTP1B, n=15-16 mice/group; (**H**) ELISA quantification of Aβ_1–42_ levels in the brain diethylamine (soluble) and formic acid (insoluble) fractions of cortical tissue APP/PS1 mice with or without DPM-1003 treatment, n=9-11 mice/group.

Considering both soluble Aβ and insoluble Aβ aggregates contribute to AD pathology (*2*, *3*), we examined whether PTP1B deletion altered levels of both forms in APP/PS1 mice. We performed sequential extraction of brain tissues using diethylamine for soluble Aβ and formic acid for insoluble Aβ (*50*). Notably, ELISA quantification revealed that PTP1B deletion in APP/PS1 mice reduced soluble Aβ_42_ levels by 48% and insoluble Aβ_42_ levels by 35% (Fig. 2G). Similarly, treatment with DPM-1003 in APP/PS1 mice resulted in reduction in soluble Aβ_42_ by 28% and insoluble Aβ_42_ by 30% (Fig. 2H). These findings suggest that both genetic deletion and pharmacological inhibition of PTP1B attenuate amyloid pathology in APP/PS1 mice.

The levels of Aβ in the brain are dependent on the dynamic equilibrium between Aβ production and clearance. Aβ is produced through the proteolytic processing of APP, by both β- and γ-secretases. Interestingly, immunoblot analysis of brain lysates of APP/PS1 mice revealed that whereas PTP1B deletion reduced Aβ levels, it did not alter the expression levels of full-length APP, APP C-terminal fragments (APP-CTFs), or key APP-processing enzymes, including Beta-site APP cleaving enzyme-1 (Bace-1) and Presenilin 1 (PS1), a component of the γ-secretase complex (fig. S3B-S3C). Notably, PS1 expression was reduced in APP/PS1 mice due to the presence of a PS1 deletion mutation. These findings suggest that the reduction in Aβ levels following PTP1B deletion is likely driven by enhanced Aβ clearance rather than altered APP expression or processing in APP/PS1 mice.

### PTP1B is highly expressed in brain immune cells and limits microglial reactivity in APP/PS1 mice

In light of the reduced Aβ burden in APP/PS1;PTP1B^-/-^ mice, and since microglia are known to play a crucial role in Aβ uptake and degradation, we hypothesized that these effects may be driven by enhanced microglial activity. To investigate this, we conducted an unbiased assessment of gene expression changes in the brains of APP/PS1 mice using single-cell RNA sequencing. Specifically, we used 8 age-matched (13–month-old) female mice, equally divided between APP/PS1, and APP/PS1; PTP1B^-/-^. Cells from the hippocampal region from 2 mice of the same genotype were pooled, yielding 4 samples for single-cell RNA sequencing. (Fig. 3A).

**Fig. 3.**
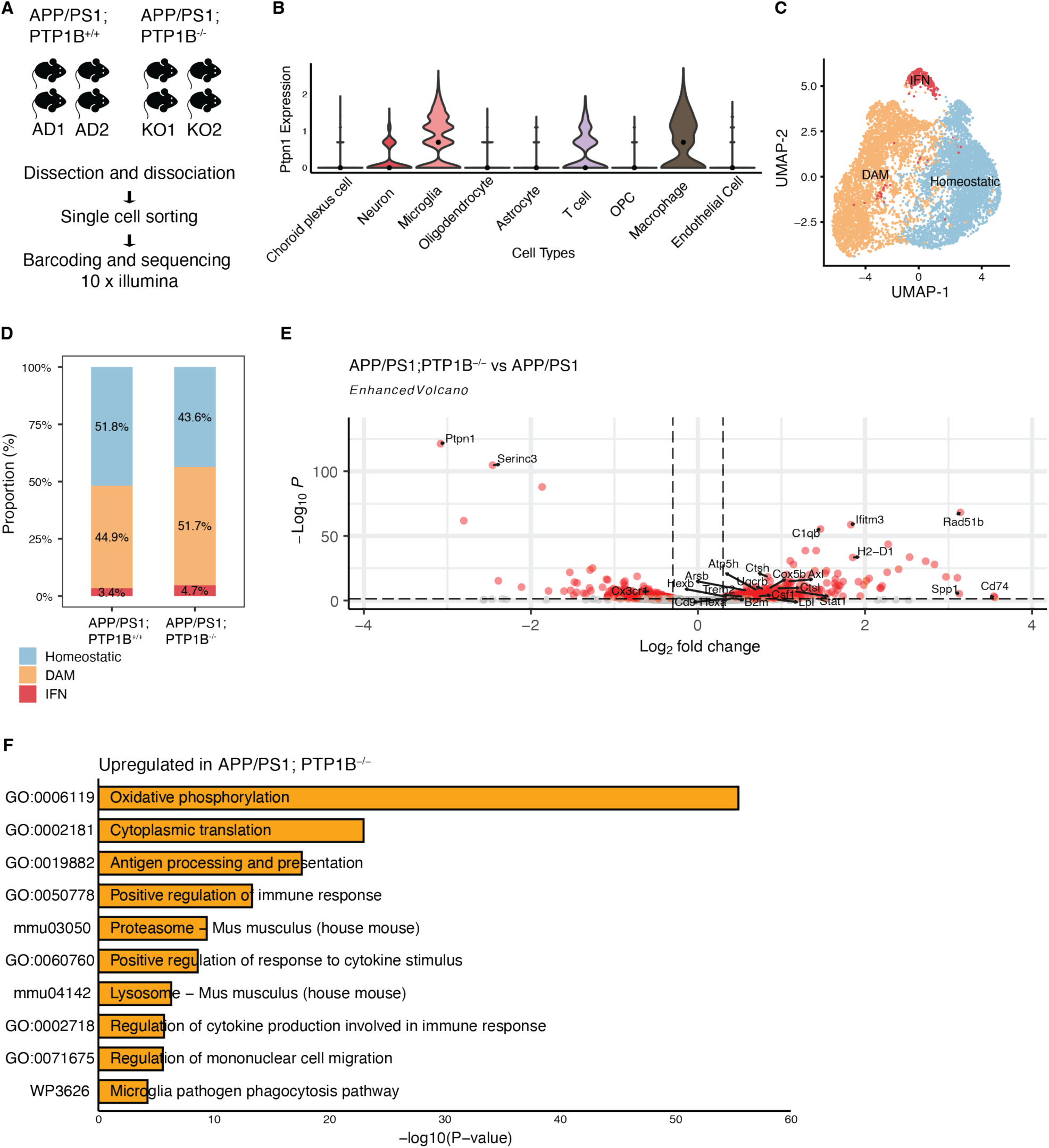
PTP1B is highly expressed in brain immune cells and limits microglial reactivity in APP/PS1 mice. (**A**) Schematic of the scRNA-seq experimental design on 13-month-old female mice. (**B**) *Ptpn1* expression across all annotated cell types in APP/PS1 mice. (**C**) UMAP plots of microglia subclusters split by genotype and colored according to subclusters. (**D**) Proportion of microglia subclusters from each genotype. (**E**) Volcano plot showing significant differential expressed genes in red dots (adjusted P < 0.05, log_2_FC> 0.3 or log_2_FC< −0.3) in microglia of APP/PS1;PTP1B^-/-^ versus APP/PS1; pseudobulk expression was generated by summing all counts per gene. (**F**) Gene ontology bar graph generated from genes significantly upregulated (adjusted P < 0.05, basemean>200, log_2_FC>0.3) in APP/PS1;PTP1B^-/-^ mice.

After quality control and filtering, a total of 31740 cells across two genotypes were arranged by uniform manifold approximation and projection (UMAP) for visualization (fig. S4A). Clusters were then manually annotated as astrocytes, choroid plexus cells, endothelial cells, macrophages, microglia, neurons, oligodendrocytes, OPCs and T cells, based on the expression of cell type-specific signature genes (fig. S4B). Then, we investigated the expression level of PTP1B across all cell types in the brain. Interestingly, *Ptpn1*, the gene that encodes PTP1B, was found to be highly expressed in immune cells, including microglia, macrophages, and T cells (Fig. 3B), suggesting a potential role in brain immune regulation.

As microglia are the primary phagocytes in the brain, we subclustered them for further analysis. This revealed three distinct subclusters: homeostatic microglia (Homeostatic; expressing markers: *Fcrls*, *Tmem119* and *P2ry12*), disease-associated microglia (DAM; expressing markers: *Axl*, *Ctsl* and *Trem2*) and interferon-responsive microglia (IFN; expressing markers: *Oasl2*, *Stat1* and *Irf7*) (fig. S3C). The proportion of DAM microglia showed a modest increase in APP/PS1; PTP1B^-/-^ mice (51.7%) compared to the controls APP/PS1 mice (44.9%), suggesting a possible shift towards a more enhanced microglial activation in the absence of PTP1B.

To examine further how PTP1B affects the microglial response to Aβ pathology, we analyzed the differentially expressed genes in the microglia. Genes upregulated in the absence of PTP1B include those encoding major histocompatibility complex (MHC) proteins (e.g. *H2-D1*, *Cd74* and *B2m*), interferon response proteins (e.g. *Stat1*, *Ifitm3* and *Oasl2*), members of the cathepsin family of cysteine proteases (e.g. *Ctsh*, *Ctsl* and *Ctsc*) and lysosomal proteins (e.g. *Hexa*, *Hexb* and *Arsb*). Additionally, several genes associated with DAM signatures were upregulated, including *Axl*, *Cd9*, *Trem2*, *Csf1* and *Spp1*. Some microglial homeostatic genes, such as *Serincs* and *Cx3cr1*, were downregulated in PTP1B-deficient microglia (Fig. 3E). These findings revealed that PTP1B deletion in APP/PS1 mice drives a transcriptional shift towards a more active and phagocytic state. Indeed, gene ontology (GO) pathway analysis showed enrichment for processes involving the “proteosome”, “lysosome” and “microglia pathogen phagocytosis pathway” (Fig. 3F). Interestingly, PTP1B-deficient microglia exhibited an enrichment of genes related to oxidative phosphorylation (OXPHOS) pathway. This metabolic enhancement likely supports the increased activation and phagocytosis. Together, these data suggest that microglia could be more activated and phagocytic in APP/PS1 mice upon the deletion of PTP1B.

### PTP1B deficiency enhances microglial phagocytic activity in response to Aβ

Considering the reduction of Aβ levels (Fig. 2) and microglial transcriptional changes (Fig. 3) observed in APP/PS1 mice lacking PTP1B, we hypothesized that PTP1B deletion would increase microglial phagocytic activity. To test this, an assay measuring uptake of fluorescent beads was performed on primary microglia isolated from WT and PTP1B-knockout pups. Phagocytic activity was quantified by flow cytometry as the percentage of cells that internalized the fluorescent beads. In WT microglia, treatment with Aβ oligomers (AβOs) increased phagocytic activity compared to vehicle-treated control. Interestingly, PTP1B-deficient microglia exhibited a higher baseline phagocytic activity, and this effect was further enhanced upon stimulation with AβOs (Fig. 4A and fig. S5A). These data suggest that PTP1B deletion enhances microglial phagocytic capacity in vitro, supporting a role of PTP1B as a negative regulator of microglial phagocytosis.

**Fig. 4.**
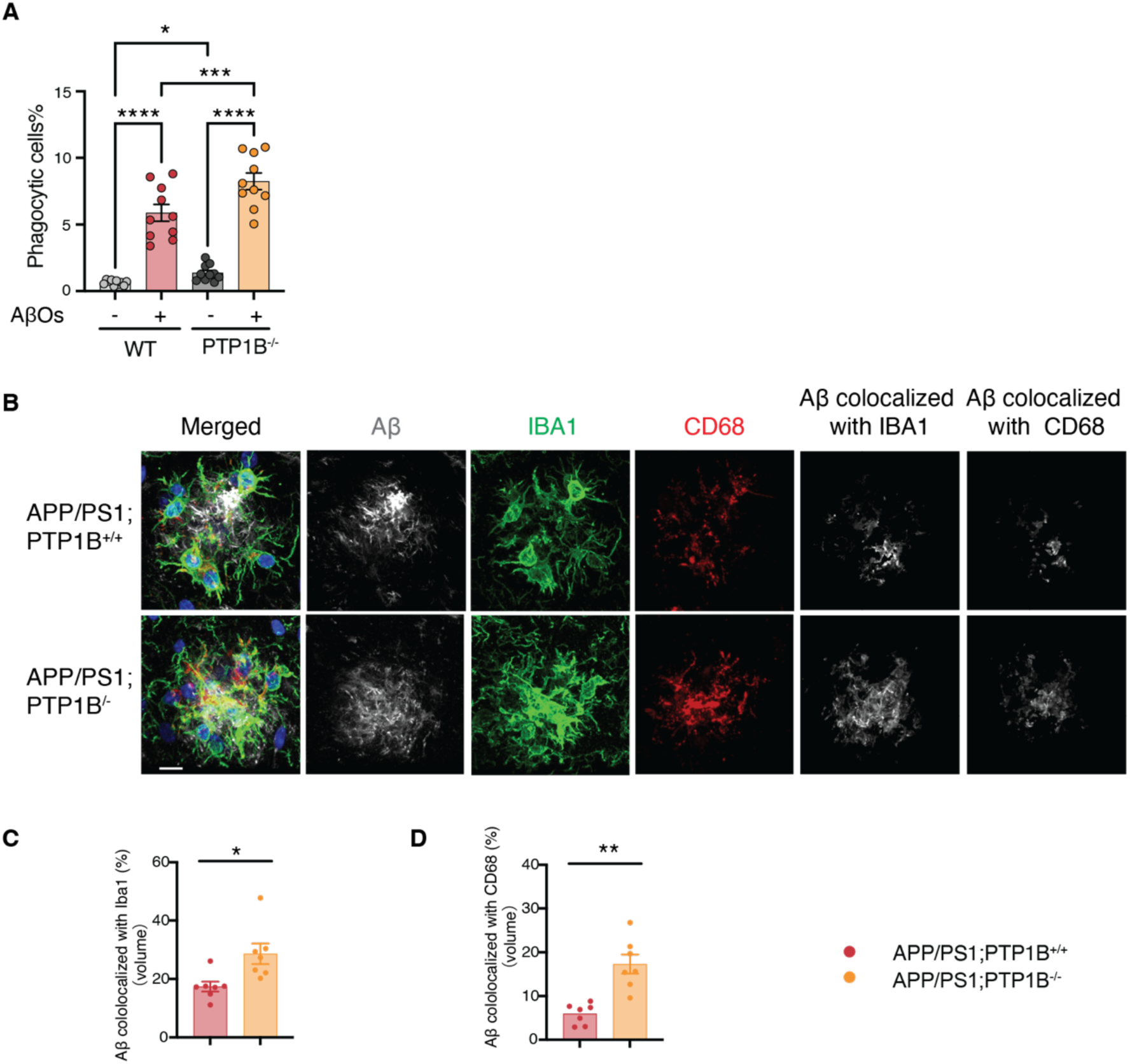
PTP1B deletion enhanced Aβ engulfment in APP/PS1 mice and promoted phagocytosis in vitro. (A) Quantification of WT and PTP1B^-/-^ primary microglia, treated with vehicle or AβOs, and measurement of phagocytic activity by fluorescence-activated cell sorting (FACS)-based microparticle-uptake assay (n = 9 per group). (B) Representative images of microglia (IBA1, pseudo green), Aβ plaques (6E10, pseudo grey) and lysosomal marker (CD68, pseudo red) co-stained in APP/PS1 mice with or without PTP1B; nuclei are stained with DAPI (blue); Aβ engulfed by IBA1^+^ microglia and by CD68^+^ microglia were shown in grey, scale bar=10µm. (**C**) Quantification of the percent volume of Aβ engulfed by IBA1^+^ microglia, averaged across multiple images per mouse; each dot represents one mouse (n=7 mice per group). (**D**) Quantification of the percent volume of Aβ engulfed by CD68^+^ microglia, averaged across multiple images per mouse; each dot represents one mouse (n=7 mice per group).

To assess whether this effect was recapitulated in vivo, we performed immunohistochemistry and quantitative confocal imaging on brain slices from APP/PS1 mice, in the presence or absence of PTP1B (Fig. 4C and fig. S5B). We quantified Aβ engulfment by calculating the volume of Aβ signal colocalized with IBA1-positive microglia, normalized to the total Aβ volume. Notably, PTP1B-deficient microglia exhibited a significant increase in the percentage of intracellular Aβ compared to controls (Fig. 4C). Additionally, immunofluorescence staining of CD68, a lysosomal marker of phagocytic microglia, revealed that Aβ internalization within CD68^+^ phagolysosomes was more than doubled in the absence of PTP1B (Fig. 4B and 4D). These results suggest that enhanced Aβ plaque engulfment by PTP1B-deficient microglia may contribute to the reduced Aβ burden observed in APP/PS1 mice lacking PTP1B.

### PTP1B deletion enhanced AβO-induced PI3K-AKT-mTOR signaling and energy metabolism in microglia

The activation of myeloid cells, such as microglia, is regulated by the PI3K-AKT-mTOR signaling pathway, which plays a critical role in cellular metabolism (*37*, *51*). Upon activation, this pathway promotes the phosphorylation of AKT and mTOR, leading to the upregulation of HIF1α, a transcriptional regulator of glycolysis. Enhanced glycolysis enables microglia to generate ATP rapidly to meet the immediate energy demands associated with immune functions such as phagocytosis. Although glycolysis provides a fast and flexible energy source, OXPHOS remains a more efficient pathway of sustaining longer-term energy production. Both pathways can contribute to the metabolic demands of activated microglial function (*36–38*, *40*).

To investigate the role of PTP1B in this metabolic regulation, we examined signaling changes in primary microglia with or without PTP1B following exposure to AβOs. In WT microglia, stimulation with AβOs increased phosphorylation of AKT and mTOR and elevated HIF1α expression. Interestingly, PTP1B-knockout microglia displayed further enhancement of the PI3K-AKT-mTOR pathway, as demonstrated by increased AKT and mTOR phosphorylation, and increased HIF1α level (Fig. 5A), suggesting that PTP1B deletion amplifies microglial signaling responses to AβOs.

**Fig. 5.**
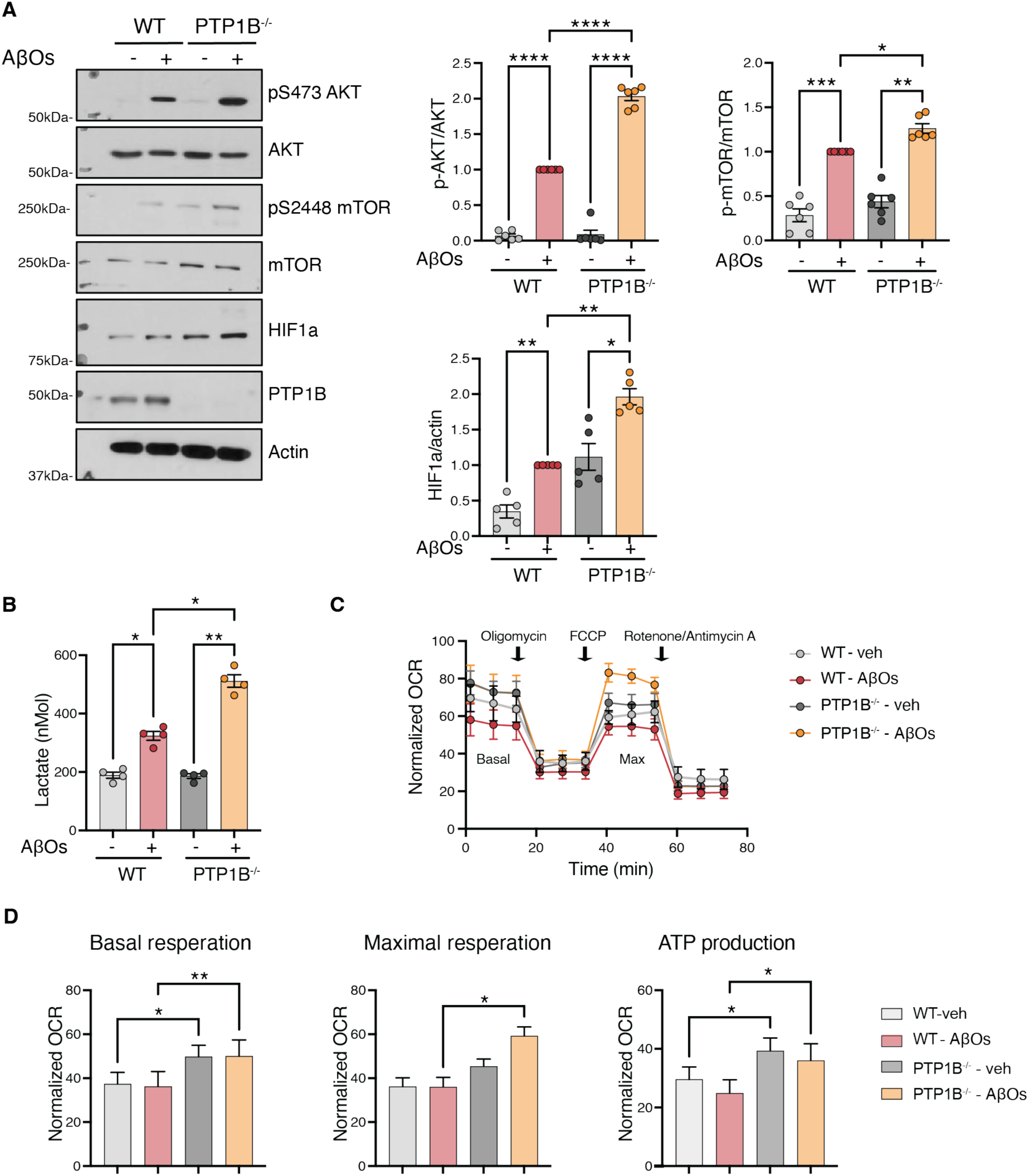
PTP1B deletion enhanced AβOs-induced PI3K-AKT-mTOR signaling and energy metabolism in microglia. (**A**) Immunoblot analysis and quantification of AKT, p-AKT, mTOR, p-mTOR, HIF1α, and Actin in WT or PTP1B^-/-^ primary microglia treated with vehicle or AβOs. (**B**) Glycolysis measured by lactate secretion (n = 4 per group). **(C-D**) WT and PTP1B^-/-^ primary microglia were treated with and without AβOs for 24h. After treatment, real time oxygen consumption rate (OCR) was measured following sequential addition of oligomycin, FCCP, rotenone and antimycin A as illustrated (n = 3 independent experiments, with 5-8 technical replicates per experiment); (C) Oxygen consumption rate; (D) Quantification of basal respiration, maximal respiration and mitochondrial-linked ATP production; Oligomycin: an ATP synthase inhibitor; FCCP: a proton ionophore; rotenone and antimycin A: electron transport chain inhibitors.

To assess downstream metabolic effects, we measured both glycolytic output and mitochondrial respiration of WT and PTP1B-deficient microglia. Glycolytic activity was first evaluated by measuring lactate secretion into the culture medium, as lactate is a by-product of glycolysis. Stimulation with AβOs increased lactate production in WT microglia, whereas in PTP1B-deficient microglia lactate production was further increased, suggesting elevated glycolytic metabolism in the absence of PTP1B (Fig. 5B). Consistently, Seahorse analysis revealed a higher extracellular acidification rate (ECAR) in PTP1B-deficient microglia in response to AβOs (fig. S6), indicative of increased glycolytic flux. In the same assay, measurement of the oxygen consumption rate (OCR) illustrated that PTP1B deletion also led to an increase in mitochondrial respiration, as indicated by elevated basal respiration under both vehicle and AβO-treatment conditions (Fig. 5C-D). Additionally, PTP1B-deficient microglia exhibited enhanced maximal respiration upon stimulation with AβOs (Fig. 5D), indicating a greater respiration capacity. Similarly, mitochondrial-linked ATP production was higher in PTP1B-deficient microglia under both vehicle and AβO-treatment conditions (Fig. 5E). Together, these findings suggest that PTP1B deletion in microglia enhances both mitochondrial and glycolytic metabolism, promoting a metabolically active state that may support enhanced microglial function such as phagocytosis.

### PTP1B plays a critical role in regulating microglia activation upon AβOs stimulation via SYK-dependent pathway

Since we demonstrated that PTP1B deletion enhanced phagocytic activity (Fig. 4) and energy metabolism (Fig. 5) in microglia in response to AβOs, we examined upstream signaling pathways to identify mediators of these responses. We focused on SYK, a protein tyrosine kinase previously reported to be a central node in the microglial responses to Aβ pathology (*41–43*). In addition, SYK was reported to act upstream of the PI3K-AKT-mTOR signaling pathway, playing a key role maintaining microglia fitness (*41–43*).

Interestingly, following treatment with AβOs, PTP1B^-/-^ microglia exhibited increased SYK phosphorylation compared to WT (Fig. 6A), suggesting PTP1B acts as a negative regulator of SYK. To determine whether SYK signaling contributes to the role of PTP1B in microglial activation, we pretreated both WT and PTP1B^-/-^ microglia with the SYK inhibitor BAY61-3606 at varying concentrations. Treatment with BAY61-3606 effectively reduced SYK tyrosine phosphorylation in both WT and PTP1B^-/-^ microglia, indicating successful inhibition of SYK activity (Fig. 6B). Functionally, the microparticle uptake assay showed that treatment with BAY61-3606 diminished the enhanced phagocytic activity of PTP1B^-/-^ microglia, reducing it to the levels seen in treated WT microglia (Fig. 6C and fig. S7A). Immunoblot analysis further demonstrated that BAY61-3606-treated microglia displayed reduced AKT-mTOR signaling in both groups (Fig. 6B and fig. S7B). Consistently, treatment with SYK inhibitor led to a reduction in lactate production and basal oxygen consumption rate to levels similar to both WT and PTP1B-deficient microglia (Fig. 6D-E). Taken together, these data demonstrate that PTP1B can regulate SYK function, which in turn plays an important regulatory role in controlling microglial responses to AβOs, including phagocytosis, signaling and metabolic fitness.

**Fig. 6.**
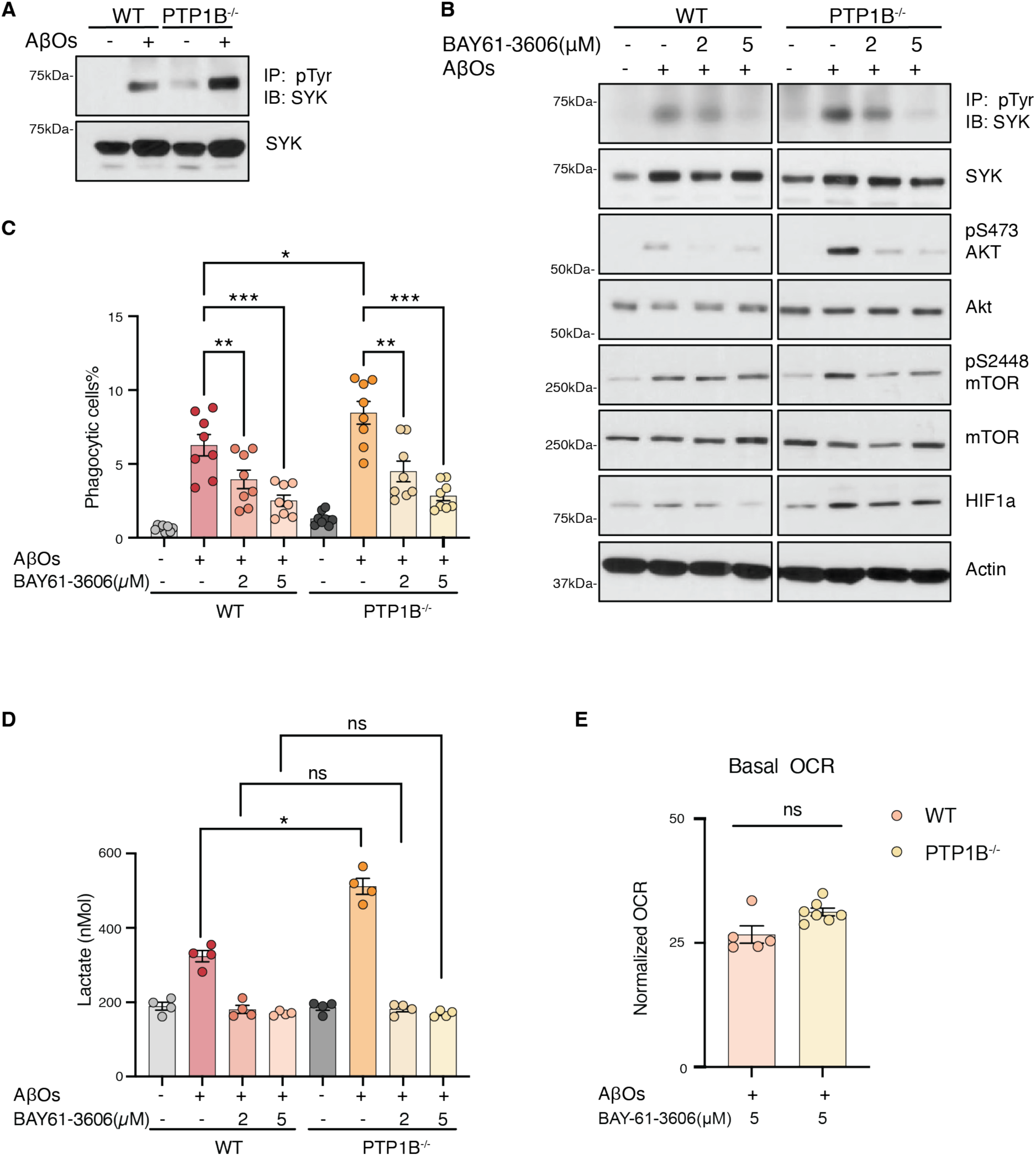
PTP1B plays a critical role in regulating microglia activation upon AβOs stimulation via SYK-dependent pathway. (**A**) Representative immunoblot analysis of SYK and p**-**SYK in WT or PTP1B^-/-^ microglia treated with vehicle or AβOs; p-SYK was detected by immunoprecipitation with anti-phosphotyrosine antibodies followed by immunoblotting for total SYK; cell lysates were prepared using RIPA buffer. (**B**) Representative immunoblot analysis of SYK, p**-**SYK, AKT, p-AKT, mTOR, p-mTOR, HIF1α, and Actin in WT or PTP1B^-/-^ microglia treated with AβOs or varying concentrations of SYK inhibitor BAY-61-3606. (**C**) Quantification of microglia and measurement of phagocytic activity by fluorescence-activated cell sorting (FACS)-based microparticle-uptake assay (n = 7**/**group). (**D**) Glycolysis measured by lactate secretion (n = 4**/**group). (**E**) OCR measurements of primary cultured microglia in the presence of BAY61-3606.

### SYK is a direct PTP1B substrate

A previous study in B cells suggested that SYK signaling may be regulated by PTP1B, as phosphorylation at Y525/Y526 was increased in the absence of PTP1B (*44*). However, it remains unclear whether PTP1B regulates SYK activation directly by targeting phosphotyrosine residues on SYK or indirectly through other signaling molecules. Our earlier structural studies identified a sequence motif —D/E–pY–pY–R/K— that is critical for optimal recognition of substrates by PTP1B (*52*). Notably, SYK’s activation loop features a similar motif: D-E-N-pY-pY-K. With this in mind, we hypothesized that SYK may be a direct substrate of PTP1B.

Upon overexpression in HEK293T cells (HEK293T-SYK), we observed SYK autophosphorylation (Fig.7A). Consistent with our findings in microglia, SYK phosphorylation was increased in PTP1B-KO 293T cells compared to WT controls (Fig.7A, lane 1 and 2). In examining whether PTP1B directly affected kinase activation, we observed a dose-response relationship with increasing levels of PTP1B ectopically expressed in PTP1B-deficient HEK 293T cells (Fig.7A, lane 3-7), coinciding with decreased phosphorylation of Y525/526 from the activation loop of SYK.

**Fig. 7.**
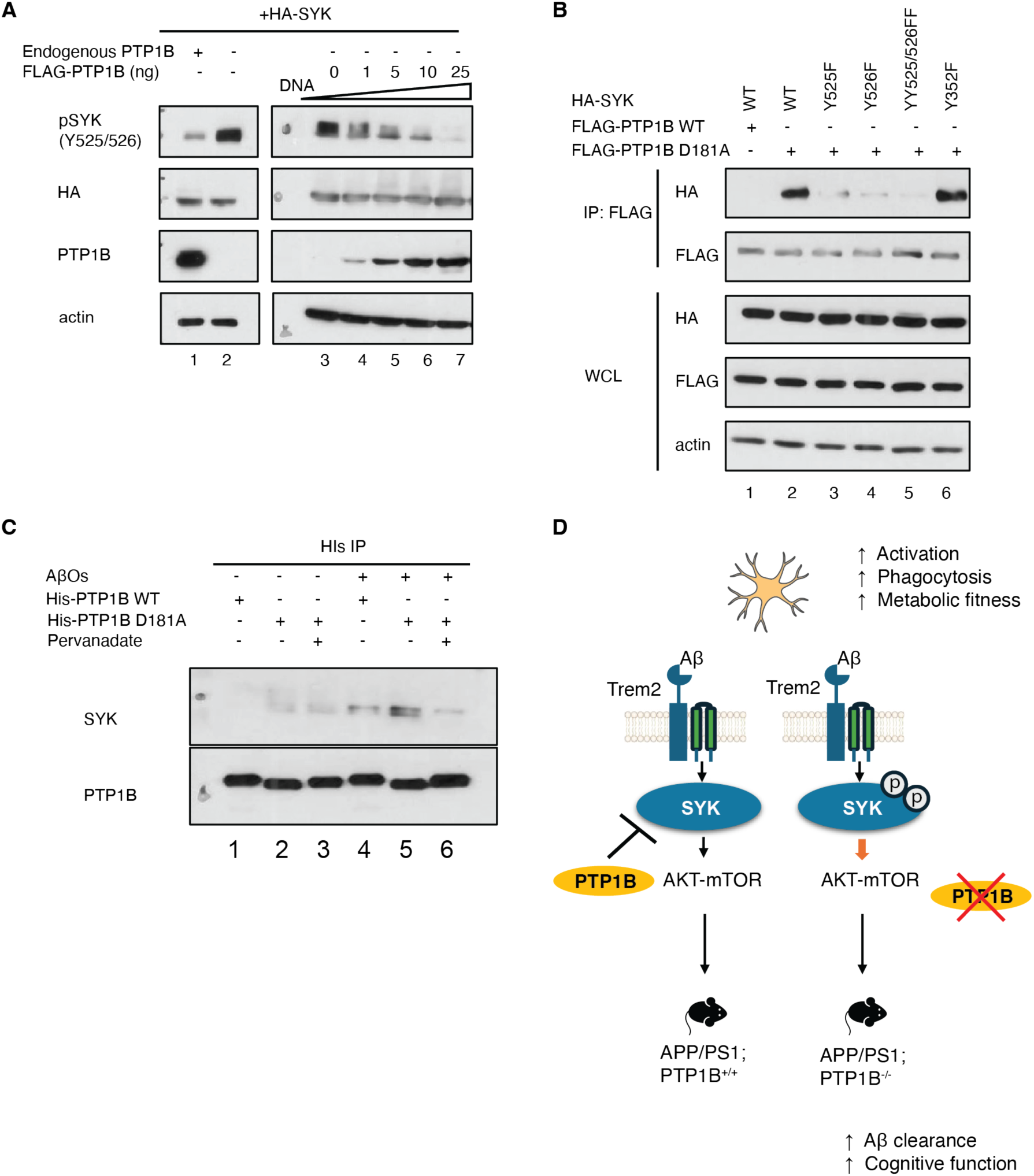
SYK is a direct substrate of PTP1B. **(A)** Immunoblot analysis of SYK phosphorylation at Y525/526 in 293T cells; the first two lanes compares SYK phosphorylation in WT and CRISPR-generated PTP1B-KO 293T cells expressing SYK; the remaining lanes show SYK phosphorylation in PTP1B-KO 293T cells co-expressing SYK and with increasing amounts of PTP1B. (**B**) Immunoblot analysis of HA-SYK co-immunoprecipitated with FLAG-PTP1B in 293T cells transiently transfected with FLAG-PTP1B (WT or D181A mutant) and HA-SYK (WT, Y525F, Y526F, YY525/526FF, or Y352F mutant) constructs, alone or in combination as indicated. Whole-cell lysates were probed for HA and FLAG**. (C**) Immunoblot of SYK and PTP1B following immunoprecipitation with purified His-tagged PTP1B (WT or D181A mutant) incubated with vehicle or AβOs -stimulated primary microglial lysates, in the presence or absence of pervanadate. (**D**) Proposed model: In microglia, PTP1B directly dephosphorylate SYK, suppressing the downstream AKT-mTOR signaling; deletion of PTP1B prevents SYK dephosphorylation, leading to enhanced AKT-mTOR signaling, which promotes increased activation, phagocytosis and metabolic fitness in response to Aβ. In APP/PS1 mice, PTP1B deletion facilitates Aβ clearance and improve cognitive function.

Additionally, we took advantage of the strategy involving substrate-trapping mutant forms of the phosphatase (*53*) to investigate whether SYK is a direct substrate of PTP1B. Unlike WT PTP1B, which dephosphorylates and releases its substrate, the PTP1B-D181A mutant forms a stable complex with its substrate, allowing for substrate identification and characterization (*53*, *54*). The substrate-trapping mutant, but not the WT form of the phosphatase, formed a stable complex with SYK (Fig.7B, lane 1 and 2), supporting a direct enzyme-substrate interaction. To identify the critical tyrosine residues mediating the PTP1B-SYK interaction, we expressed WT and mutant (Y525F, Y526F, YY525/Y526FF, Y352F) forms of SYK together with the PTP1B-D181A mutant in HEK293T cells. Mutation of the activation loop residues Y525 and Y526 in SYK to phenylalanine disrupted this interaction (Fig.7B, lanes 3-5), whereas mutation at Y352 had no observable effect (Fig.7B, lane 6). Taken together, these findings demonstrate a direct interaction between PTP1B and SYK at the critical autophosphorylation sites, Y525/Y526.

Furthermore, we tested whether PTP1B directly interacts with phosphorylated SYK in primary microglia following treatment with AβOs. Lysates of primary microglia, treated with or without AβOs, were collected and incubated with either WT or D181A mutant PTP1B protein. In the presence of AβOs, although a low level of co-precipitation of SYK with WT PTP1B was detected (Fig. 7C, lane 3), this association was more pronounced with the PTP1B-D181A substrate–trapping mutant (Fig.7C, lane 4 and 5). Furthermore, pretreatment of the PTP1B substrate trapping mutant with pervanadate, which disrupts the enzyme’s active site, prevented its binding to SYK (Fig.7C, lane 6), indicating that the interaction between PTP1B and SYK requires the PTP1B catalytic site. Therefore, these results suggest that SYK is a direct substrate of PTP1B, dephosphorylation of which represents a potential mechanism underlying the effects of PTP1B deletion or inhibition in AD mouse model (Fig 7D). Together, these findings established that PTP1B as a critical regulator in microglial function and provides new insights into its potential as a therapeutic strategy for AD.

## Discussion

Accumulating evidence suggests that metabolic alterations in the brain, such as insulin resistance and impaired glucose metabolism, are early features of AD (*10*, *55–60*). PTP1B is a widely expressed phosphatase best known for its role in metabolic regulation, particularly through its regulation of insulin and leptin signaling, both of which have been associated with neuroprotective functions (*61*). These connections led us to hypothesize that targeting PTP1B may restore the disrupted brain energy metabolism and improve cognitive outcome in AD.

PTP1B modulates diverse signaling pathways across several tissues – including insulin signaling in metabolic organs, HER2 signaling in breast cancer, and JAK/STAT cytokine signaling in immune cells. As a result, PTP1B has been explored as a therapeutic target in a range of diseases and contexts, including diabetes (*17–19*), breast cancer (*62*) and immunotherapy involving T cell checkpoint blockade (*30*). Although one study reported that lifetime deletion of PTP1B in myeloid lineage cells predisposed aged mice to leukemia (*63*), the heterozygous animals appeared normal, suggesting that partial inhibition with a PTP1B inhibitor could provide therapeutic efficacy. In the central nervous system (CNS), most studies of PTP1B have focused its role on neurons, where it negatively regulates key signaling pathways involved in neuronal survival, synaptic function, and metabolism (*24*, *64–66*). For example, PTP1B dephosphorylates the TrkB receptor in neurons, attenuating BDNF signaling (*24*). In the hAPP-J20 AD model, neuron-specific PTP1B deletion enhanced the phosphorylation of synaptic proteins such as NSF and NMDAR GluN2B, leading to improved synaptic plasticity (*65*). Thus, neuronal PTP1B has been proposed as a potential therapeutic target for neurological diseases.

Although prior studies have primarily focus on the role of PTP1B in neurons, our RNA sequencing data showed higher mRNA levels of PTP1B in microglia compared to neurons (Fig. 3B). However, its function in microglia, especially in the context of AD, remains largely unknown. Our findings revealed a novel role for PTP1B in microglia, highlighting a broad involvement in the inflammatory response within CNS. Specifically, PTP1B deletion in APP/PS1 mice reduced brain amyloid levels. This was accompanied by enhanced Aβ engulfment by microglia, suggesting that increased phagocytic activity may contribute to the observed reduction of Aβ burden. Consistent with this, single-cell RNA sequencing revealed that PTP1B is highly expressed in microglia, and its deletion in APP/PS1 mice induces a transcriptional shift toward a more metabolically active and phagocytic microglial state. These findings extend the role of PTP1B beyond neurons, highlighting its influence on microglial signaling pathways.

In AD, microglia change from homeostatic to a disease-associated phenotype, often referred to disease-associated microglia (DAM) (*67*), which is characterized by the upregulation of genes involved in Aβ recognition and clearance. Accumulating evidence suggests that promoting expression of DAM genes can enhance microglial function and may represent a promising therapeutic strategy for halting AD progression. For example, transcriptomic studies in AD models and patients treated with anti-amyloid antibodies have shown increased expression of DAM genes, suggesting that DAM activation may be a key feature of effective Aβ-targeting therapies (*34*, *35*). Similarly, activating the microglial receptor TREM2, a critical regulator of DAM progression (*67*), using agonist antibodies, has been reported to enhance microglial responses to Aβ, leading to increased microglial proliferation and metabolism (*40–42*, *68–73*). Additionally, modulating signaling molecules downstream of TREM2, such as SYK (*41–43*) and mTOR (*39*), has been shown to attenuate Aβ accumulation and promote phagocytosis, suggesting that targeting these pathways holds promise for developing novel AD therapies. Our data are consistent with these findings, showing that PTP1B deletion induces a DAM-like phenotype, enhancing Aβ clearance.

Mechanistically, we show that PTP1B directly regulates the autophosphorylation sites of SYK within its kinase domain, acting as a brake on SYK function. The absence of PTP1B leads to enhanced SYK activation, which is critical for microglial activation. Consistent with this, inhibition SYK using BAY61-3606 led to a reduction in phagocytosis activity, regardless of PTP1B presence. Since SYK signaling acts upstream of PI3K-AKT-mTOR signaling in microglia (*41*, *42*), treatment with BAY61-3606 also attenuated AKT-mTOR phosphorylation, irrespective of PTP1B presence, underscoring SYK as a key regulatory node. Together, we demonstrated that regulation of SYK by PTP1B plays a critical role in regulating microglial activation upon Aβ stimulation.

The PI3K-AKT-mTOR pathway is a key regulator of energy balance and metabolism. Microglia rely on glycolysis and OXPHOS to fulfill their energy demands and dynamically adjust these metabolic pathways upon activation (*37*). Notably, microglia lacking critical activation molecules such as TREM2 (*74*) or SYK (*42*) have impaired metabolism, reducing both glycolysis and OXPHOS, together with reduced PI3K-AKT-mTOR signaling. Interestingly, we found that PTP1B-deficient microglia increased activation of the PI3K-AKT-mTOR pathway, exhibited enhanced glycolysis and OXPHOS, particularly upon Aβ stimulation. Transcriptomic analysis further supports an increase in oxidative metabolism, with upregulation of OXPHOS-related genes in PTP1B-deficient microglia. Collectively, these results position PTP1B as a key regulator of microglial metabolism, offering novel insight into how metabolic pathways intersect with immune activation in AD. Furthermore, previous studies demonstrated that the myeloid-specific deletion of PTP1B significantly improved glucose homeostasis in insulin resistance models (*75*, *76*), suggesting that PTP1B may also modulate glucose uptake and metabolism in microglia. Targeting PTP1B could potentially enhance microglial metabolic capacity and improve glucose availability in the brain, thereby benefiting the AD patients that display brain glucose hypometabolism (*77*).

In summary, we have now established a critical signaling function of PTP1B in microglia, in addition to its reported effects in neurons (*64–66*). We identify PTP1B as a central regulator of microglial activation, metabolism, and Aβ clearance. By directly modulating SYK signaling and downstream PI3K-AKT-mTOR axis, PTP1B deletion enhances microglial phagocytosis and promotes a more metabolically active, DAM-like phenotype. These findings provide mechanistic insight into how targeting PTP1B could enhance innate immune responses and improve microglial energy metabolism in AD. Our work demonstrates that targeting PTP1B could represent a promising strategy for mitigating cognitive decline in AD patients.

### Limitations of Study

Our study revealed a new perspective on the function of PTP1B in microglia, highlighting its role in regulating immune signaling and cellular metabolism, which aligns with emerging concepts underscore the importance of neuroinflammation in the context of AD. Going forward, it will be important to generate animals in which PTP1B is specifically ablated in microglia and neurons, so we can assess directly the cell-type specific contributions to the overall disease phenotype.

## Materials and Methods

### Mice

All animal experimental procedures were performed in accordance with the approved animal care and use protocols. The institutional animal care and use committee (IACUC) at Cold Spring Harbor Laboratory (protocol number 15-6). APP/PS1 mice (stock 004462) and *Ptpn1*^-/-^ mice were obtained from the Jackson laboratory and were crossed to generate experimental animals. All mice were 12-13 months old at the time of the behavior assay; mice were transported from the animal room to the behavior room and allowed 30 minutes to habituate before beginning behavior test.

DPM-1003 was dissolved in sterile saline and administrated intraperitoneally at a dose of 5mg/kg body weight, twice a week for 5 weeks, starting at 11 months of age.

Primary microglia for culture were isolated from pups at postnatal day 1-3.

### Novel object recognition test

Mice were habituated in a white square chamber (42 × 42 cm × 42 cm) for 5 min per day over 2 consecutive days. The activity of mice was recorded with a video camera. The apparatus was cleaned with 70% alcohol and air-dried before the start of each trial. Each mouse was given 5 minutes to freely explore the chamber for habituation 24 hours before the test. The test had two phases: training and testing. For training, two identical objects were placed in diagonally opposite corners of the chamber 8–9 cm from the walls. A mouse was placed at the midpoint between the objects. After allowing 10 min to explore the objects, the mouse was returned to the colony. The testing was performed 1h after the acquisition. In the tests for place recognition, one of the objects was replaced with a novel object of a similar height and volume but different shape and appearance. For testing, the mouse was again placed in the chamber to explore the objects for 3 min. The amount of time spent exploring each object (nose sniffing and head orientation within <2.0 cm) was recorded scored using video tracking software (Noldus Ethovision XT). The novel object exploration percentage was computed as *Novel*% = *T_new_* × 100/(*T_new_* + *T_old_*), the old object exploration percentage was computed as *Old*% = *T_old_* × 100/(*T_new_* + *T_old_*), where *T_new_* is the time spent exploring the new or moved object, and *T_old_* is the time spent exploring the familiar object. Animals that did not explore any objects during the training session or only explore one object during the testing session were excluded.

### Morris water maze

The water maze used in this study comprised a circular tank 110 cm in diameter with a platform filled with tap water at a temperature of 23-25 °C. Different shapes were posted along the walls of the tank, which served as spatial reference cues. A camera was mounted above the maze to record the swimming traces in the water maze. During the acquisition trials, the platform was submerged 1 cm below the water surface, mice were placed into the maze facing different visual cues for each trial. Mice were allowed to search for a platform for 60 s. If a mouse failed to find the platform, it was guided to the platform and maintained on the platform for 10 s, and the latency was recorded as 60s. Four trials a day were conducted. Escape latency indicative of spatial memory acquisition, was recorded for each trial. On day 6, the platform was removed, and a probe test was conducted, followed 4 trials with visible platform. The percentage time spent in each of the four quadrants, the number of target (platform) area crossings and the mean speed were recorded by video tracking software (Noldus Ethovision XT). During the visible platform trials, the platform was placed at the center of the maze and raised 1-2 cm above the water surface, marked with a small flag for visibility. Animals that took an average of more than 30 seconds to locate the platform across the four trials were considered to have vision impairments or a lack of motivation and were excluded from the study.

### Brain sample preparation

Mice were anesthetized with isoflurane and transcardially perfused with 15 mL of PBS. Brains were dissected, with the left hemisphere fixed in 4% paraformaldehyde for 24 hours at 4 °C, while the right hemisphere was further dissected to isolate the cortices and hippocampi, which were then flash-frozen and stored at −80 °C. Fixed samples were transferred to 30% sucrose for 48-72h and then frozen in −80 °C. These brains were then sectioned at 50 μm in thickness using a vibratome (Leica) and stored in cryoprotectant (30% sucrose and 30% ethylene glycol in PBS) at −20 °C for downstream staining and imaging. The flash-frozen brains were thawed for protein extraction and mechanically homogenized in RIPA buffer containing phosphatase and protease inhibitor cocktail (Thermo Fisher Scientific #87785). The homogenates were stored at −80 °C until further analysis.

### Immunofluorescence microscopy

Brain slices were washed twice with PBS for 5 minutes each to remove cryoprotectant. Antigen retrieval was performed by heating the sections in sodium citrate buffer (10 mM sodium citrate, 0.05% Tween 20 in ddH₂O, pH 6.0) at 85°C for 20 min. The sections were blocked in 5% normal donkey serum with 0.5% Triton X-100 for 1h at room temperature, followed by overnight incubation at 4°C with primary antibodies. After incubation, the sections were washed three times with PBS for 20 minutes each and incubated with fluorescent secondary antibodies for 3 hours at room temperature. DAPI (1:10000) was used to stain the nuclei for 10 min at room temperature, followed by two washes with PBS. The slices were mounted on coverslips with Ibidi mounting medium. Images were captured using either an Echo fluorescence microscope or a Zeiss LSM 780 confocal microscope.

### Assessment of Aβ plaque burden

Sagittal brain sections were stained with the thioflavin S (0.01%) and 6E10 antibodies. Epifluorescence capturing images of hippocampal regions of APP/PS1 and APP/PS1;PTP1B^-/-^ mice were captured using the Echo microscope with identical exposure times across samples for each fluorophore at a magnification of 4×. The images were analyzed using the “Analyze Particles” function in Fiji, applying consistent thresholding, size (50 to infinity), and circularity (0.1 to 1) filters for quantification. A total of 3-4 brain sections were analyzed per mouse.

### ELISA of Aβ42

Aβ extraction was performed as previously described (*50*). Briefly, mouse cortical brain tissue was homogenized in tissue homogenate buffer (2mM Tris pH 7.4, 250mM sucrose, 0.5mM EDTA, and 0.5mM EGTA in ddH₂O) with protein phosphatase and protease inhibitor. Diethylamine (DEA, 0.4%, Sigma-Aldrich) was added to the samples at a 1:1 ratio, and the homogenates were centrifuged at 135,000 × g for 1 hour at 4 °C. The supernatants were collected for analysis of the DEA-soluble Aβ fraction. The remaining pellet was resuspended in ice-cold formic acid centrifuged at 109,000 × g for 1 hour at 4 °C. The resulting supernatants were diluted 1:20 in formic acid neutralization buffer (1 M Tris base, 0.5 M Na₂HPO₄, and 0.05% NaN₃ in ddH₂O) and collected for analysis of the insoluble Aβ fraction. The protein concentration of both the soluble and insoluble fractions was measured using the Bradford Protein Assay Kit (Thermo Fisher Scientific, #23200). Aβ_1-42_ levels in both fractions were quantified using an Amyloid beta 42 Human ELISA Kit (Thermo Fisher Scientific #KHB3441) according to manufacturer instructions and normalized to the protein concentration quantified by Bradford assay.

### Single cell RNAseq

13-month-old female mice from APP/PS1;PTP1B^+/+^ and APP/PS1;PTP1B^-/-^ groups were sacrificed for the sample preparation. Mice were perfused with perfusion buffer (cold HBSS), and hippocampal tissues were dissociated in dissection buffer (DPBS Gibco, 14-287-080). Tissues dissociation was performed using the Adult Brain Dissociation Kit (Miltenyi Biotech, 130-107-677) according to the manufacturer’s instructions, with manual trituration. Briefly, the chopped tissues were incubated in digestion buffer at 37 °C in a water bath for 15 min, gently tapping the tubes 3-4 times during the digestion. Following digestion, the tissues were triturated with fire-polished glass pipettes to obtain a single cell preparation, which was then filtered through a 70 μm cell strainer. To minimize ex vivo microglial activation, transcriptional and translational inhibitors were added in perfusion, dissection and digestion buffer as previously described (*78*). In each sample, cells from two animals were pooled together.

FACS-based viability enrichment and single-cell library preparation were carried out at the Cold Spring Harbor Laboratory Single-Cell Biology Shared Resource. Prior to FACS, Single-cell suspensions were washed by centrifugation at 4°C, 600g in a swinging bucket rotor, and resuspended in 1X PBS with 0.04% BSA, and stained with SytoxBlue dead cell stain (ThermoFisher #S34857). Viable, intact cells were sorted using a Sony SH800 sorter on a 70um chip into PBS/BSA, pelleted as above and resuspended at 1,000 cells/μL. Single-cell RNA sequencing was performed using 10X Genomics NextGEM v3.1 3’ Gene Expression kits. After sorting, cell count and viability were determined using a Countess FL II automated cell counter (ThermoFisher) and ViaStain AOPI (Nexcelom #CS2-0106-5mL) live/dead cell stain. Cells were loaded into a 10x Genomics Chromium Controller and libraries prepared according to the manufacturer’s instructions to target a recovery of ∼6,000-10,000 viable cells per sample. Unique dual-indexed libraries were paired end sequenced using the Illumina NextSeq2000 sequencing system with 28x10x10x90bp reads to a mean depth of ∼50,000 reads per cell.

### Data processing

The Cell Ranger count pipeline (v7.1.0, 10X Genomics) was used to align FASTQ reads to the mouse reference genome (gex-mm10-2020-A, 10X Genomics) and produce digital gene-cell counts matrices with default settings, which includes intronic reads when tallying expression. Empty droplet removal and ambient RNA subtraction were carried out simultaneously using Cellbender (*79*) with default settings and epochs=150.

R studio (2024.9.0.375) was used for all downstream analyses and Seurat (v.5.1.0) was used for filtering out low-quality cells, normalization of the data, determination of cluster defining markers and graphing of the data on UMAP. Cells with low quality (i.e., fewer than 500 or more than 6000 detected genes, total RNA counts outside 1,000–30,000, or more than 10% mitochondrial content) and those flagged as potential doublets by scDblFinder were excluded. Each dataset was normalized with SCTransform. Integration was performed using canonical correlation analysis (CCA) by selecting 3,000 integration features, followed by data integration with Seurat’s SCT-based methods. Dimensionality reduction was conducted with PCA and UMAP (30 PCs), and graph-based clustering identified major cell populations, which were then manually annotated based on marker gene expression. A microglial subset was further isolated, re-processed, and sub-clustered to profile specific microglial states. Pseudobulk counts were generated by aggregating cell-level data per sample and cell type using Seurat’s AggregateExpression, and differential expression in microglia was subsequently assessed with DESeq2(1.46.0). Genes exhibiting a log2 fold change greater than 0.3, basemean greater than 200, and p-value below 0.05 were used for gene ontology analysis of microglial subcluster using Metascape (http://metascape.org) (*80*).

### Preparation of AβOs

Aβ oligomers (AβOs) were prepared as previously described(*37*). Aβ_1-42_ peptides (Bachem, #4061966) were dissolved in 1,1,1,3,3,3-hexafluoro-2-propanol (HFIP; Sigma-Aldrich) to a final concentration of 1 mM. After overnight evaporation of HFIP, the dried peptides were dissolved in DMSO to 1 mM, diluted in DMEM to 10 μM, and incubated overnight at 4°C.

### Primary microglia culture and stimulation

Primary microglial cultures were prepared as previously described (*37*) with modification. Briefly, cortical and hippocampal tissues from WT or PTP1B⁻/⁻ mice (postnatal day 1–3) were dissected. After meninge removal and tissue trituration, cells were plated T-75 flasks and grown in culture medium (DMEM containing 10% FBS) for 1 week to establish mixed glial cultures. L-929 cell-conditioned medium (LCM, 4 ml) was added on days 7 and 9. Primary microglia were harvested after 10–12 days by shaking flasks (600 rpm, 2 hours) and once every 3 days thereafter (up to two harvests). Isolated microglia were plated at 1.5-2.0 × 10^5^ cells/ml for 3–4 days before use. Microglia culture medium (DMEM with 10% FBS, 1% P/S, and 10% LCM) was changed every 1–2 days. Cells were treated with vehicle (DMEM containing anhydrous DMSO) or 1 μM Aβ_1–42_ for 24h. In selected experiments, primary microglia were pre-treated with BAY-61-3606 (Selleckchem, S7006) at indicated concentrations to inhibit SYK signaling. Following a 1-hour pre-treatment, vehicle or AβOs was added to the culture.

### Microparticle uptake assay

The phagocytic activity of microglia was measured by microparticle uptake assay as previously described (*37*) with modifications. Briefly, 5.0 × 10^6^ fluoresbrite Yellow Green Carboxylate Microspheres (Polysciences, #17147-5) were opsonized in DMEM with 50% FBS at 37°C for 30 minutes, diluted 1:10 in DMEM with 5% FBS, and added to cultured primary microglia (5 × 10^5^ cells/ml) for 30 minutes at 37°C with shaking. After washing with PBS, cells were detached by cell scrapers, stained with anti-CD11b antibody, and analyzed by flow cytometry (FACSCalibur). Microsphere uptake was quantified using FlowJo software.

### Measurement of lactate levels

Lactate levels in primary microglia culture medium were measured using lactate assay kit (Abcam, #ab65331) according to the manufacturer’s instructions.

### Metabolic assays

Real-time oxygen consumption rate (OCR) and extracellular acidification rate (ECAR) was measured using a seahorse XF96 analyzer (Seahorse Bioscience) with Seahorse XF Cell Mito Stress Test Kit (Agilent, #103015-100), following the manufacturer’s instructions with minor modifications. Briefly, a total of 1.0 × 10^4^ cells were seeded on the XF96 cell culture microplate, cultured and treated as specified. The assay medium (XF base medium containing 1 mM pyruvate, 4 mM glutamine, and 25 mM glucose) was prepared immediately before the assay as previously described (*37*). ECAR and OCR were measured in response to 1.5 μM oligomycin, 2.0 μM FCCP and 0.5 μM rotenone/ antimycin A to determine mitochondrial basal respiration, proton leak, maximal respiration and ATP production linked to oxidative phosphorylation. Data were analyzed using Wave software (version 2.4.3.7) and normalized to total cell number, as determined by crystal violet staining after the assay.

### Immunoblotting

Brain lysates, primary microglia lysates and HEK293T cell lysates were resuspend in RIPA buffer with phosphatase and protease inhibitor cocktail. Protein concentration was measured by Bradford protein assay kit (Thermo Fisher Scientific #23200). 5X SDS loading dye was added to protein lysates and heated at 95 °C for 8 minutes. 15-50 μg protein was loaded per lane on either 10% or 15% SDS-polyacrylamide gels and ran at 90V for 30 minutes and then 130V for 60-90min. Proteins were transferred to nitrocellulose membrane (vwr #10600004) at 90V for 2h at 4°C. Primary antibodies were incubated overnight at 4 °C. Corresponding secondary antibodies were incubated for 1h at room temperature.

### Substrate trapping assays

Substate trapping experiments were performed as described previously(*53*). HEK293T cells were cultured in DMEM supplemented with 10% FBS at 37°C and 5% CO2 in a humidified atmosphere. For transfection, HA-tagged WT SYK, SYK mutants (SYK-Y525F, SYK-Y526F, SYK-YY525/526FF, SYK-Y352F), FLAG-tagged WT PTP1B, and PTP1B mutant (PTP1B-D181A) were used. Cells were transfected with Lipofectamine 2000 (Life Technologies) for 36 hours following the manufacturer’s instructions. Flag-tagged PTP1B was immunoprecipitated using anti-FLAG magnetic agarose (Thermo Fisher Scientific, #A36797). And the resulting complexes were analyzed by immunoblotting.

Primary microglia were cultured and treated as described, and lysates were incubated with His-tagged WT PTP1B and PTP1B-D181A protein for 2 hours with or without 1 mM pervanadate. His-tag magnetic beads (Cell signaling, 8811S) were added and incubated for an additional 2 hours. After thorough washes, complexes were analyzed by immunoblotting.

### Antibodies

The antibodies used in this study are listed in Supplementary Table 1.

### Statistical analysis

Results are presented as means ± SEM. Data were analyzed with GraphPad Prism (version 10.2.3). Student’s t tests were used for comparisons between two groups. For multiple group comparisons, one-way or two-way repeated measures analysis of variance (ANOVA) was performed, followed by Tukey’s post hoc test unless otherwise noted. For Seahorse OCR quantification, data represent three independent experiments (n = 3), each with 5–8 technical replicates per condition. Technical replicates were averaged within each experiment to yield a single value per condition. One-way ANOVA with Bonferroni-corrected multiple comparisons was performed to compare preselected pairs of conditions (WT–veh vs. PTP1B⁻/⁻–Veh; WT–AβOs vs. PTP1B⁻/⁻–AβOs). * or #: p<0.05; ** or ##: p<0.01; *** or ###: p<0.001; **** or ####: p<0.0001.

## Acknowledgments

We are very grateful to DepYmed Inc.(NY), for providing the DPM-1003 that was used in this study. This study utilized Single-Cell Biology, Flow Cytometry and Microscopy Shared Resources of the Cold Spring Harbor Laboratory Cancer Center.

## Author contributions

Conceptualization: YC, LVA, NKT

Methodology: YC, SRA, DS, JP, NKT

Investigation: YC, SRA, DS, JP

Visualization: YC

Supervision: LVA, NKT

Writing—original draft: YC, NKT

Writing—review & editing: YC, SRA, DS, LVA, NKT

