## Supplementary Materials for "PTP1B inhibition promotes microglial phagocytosis in Alzheimer’s disease models by enhancing SYK signaling"

**Fig. S1.**

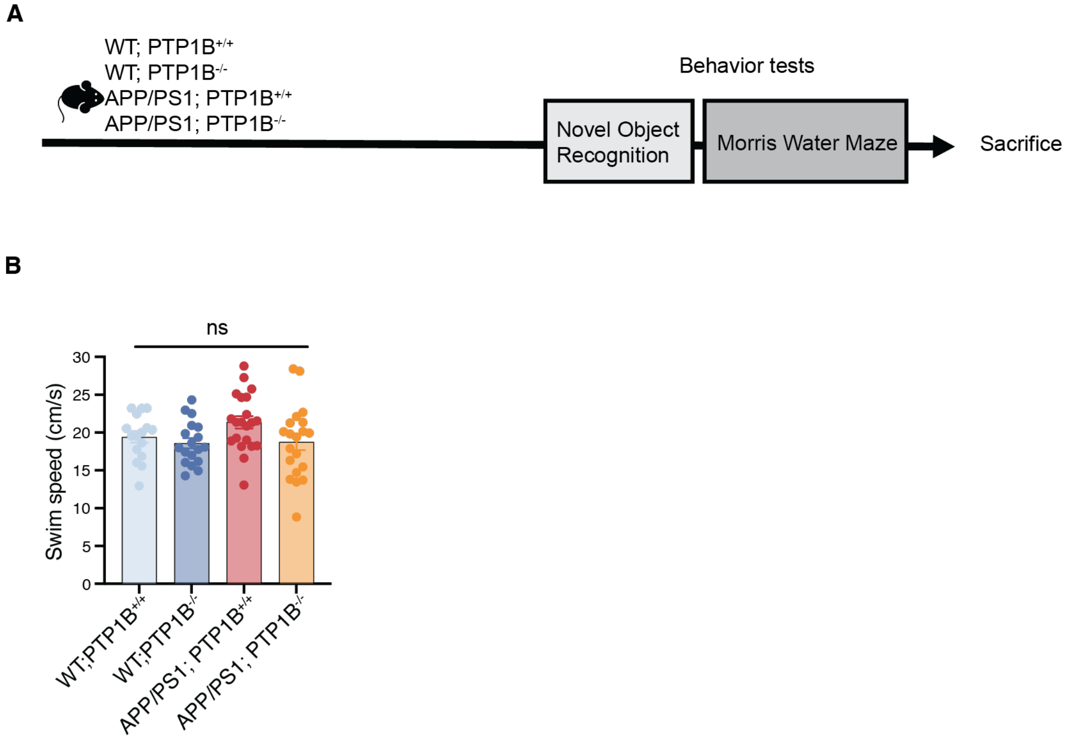


**Supplementary Figure 1 – related to Figure 1**

**(A)** Schematic of the behavioral testing timeline in 12- to 13-month-old mice with or without PTP1B deletion.

**(B)** Swimming speed during the Morris water maze probe testing on WT or APP/PS1 mice with or without PTP1B deletion.

**Fig. S2.**

A

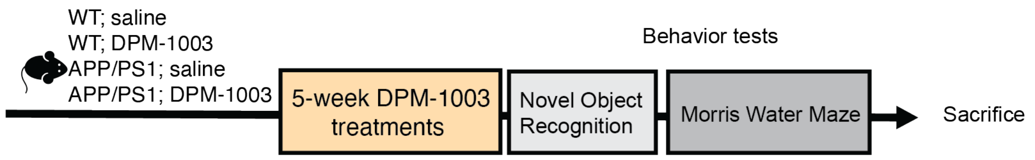


B

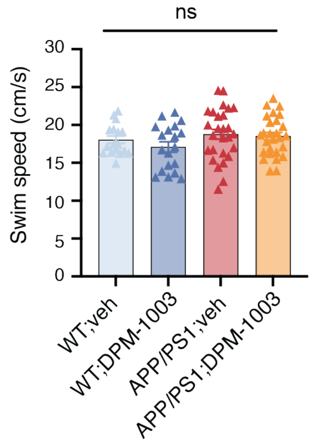


**Supplementary Figure 2 – related to Figure 1**

**(A)** Schematic of treatment in 11-month-old mice receiving vehicle or DPM-1003 treatment for 5 weeks and subsequent behavioral tests.

**(B)** Swimming speed during the Morris water maze probe testing on WT or APP/PS1 mice treated with or without DPM-1003.

**Fig. S3.**

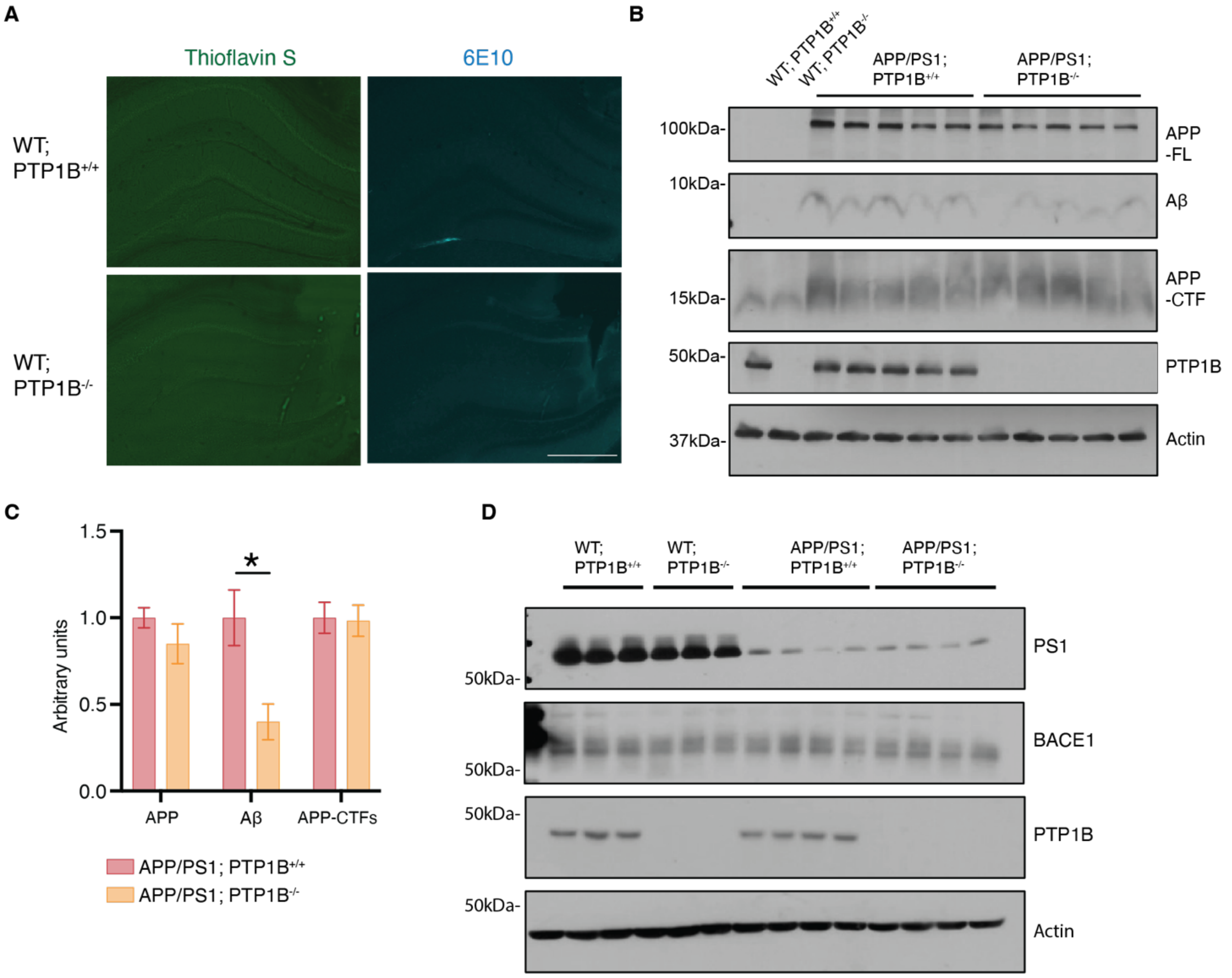


**Supplementary Figure 3 – related to Figure 2**

**(A)** Immunofluorescence of Aβ levels in WT mice with or without PTP1B in hippocampal region, Scale bar =500µm. **(B)** Immunoblot of full-length APP, Aβ, C-terminal fragments, PTP1B and Actin from 13-month-old APP/PS1;PTP1B$^{+/+}$, APP/PS1;PTP1B$^{-/-}$ and control mice brain lysates. **(C)** Quantification off full-length APP (APP-FL), APP C-terminal fragments (APP-CTFs) and Aβ normalized to Actin; **(D)** Immunoblot of Presenilin-1, BACE1, PTP1B and Actin from 13-month-old APP/PS1;PTP1B$^{+/+}$, APP/PS1;PTP1B$^{-/-}$ and control mice brain lysates.

**Fig. S4.**

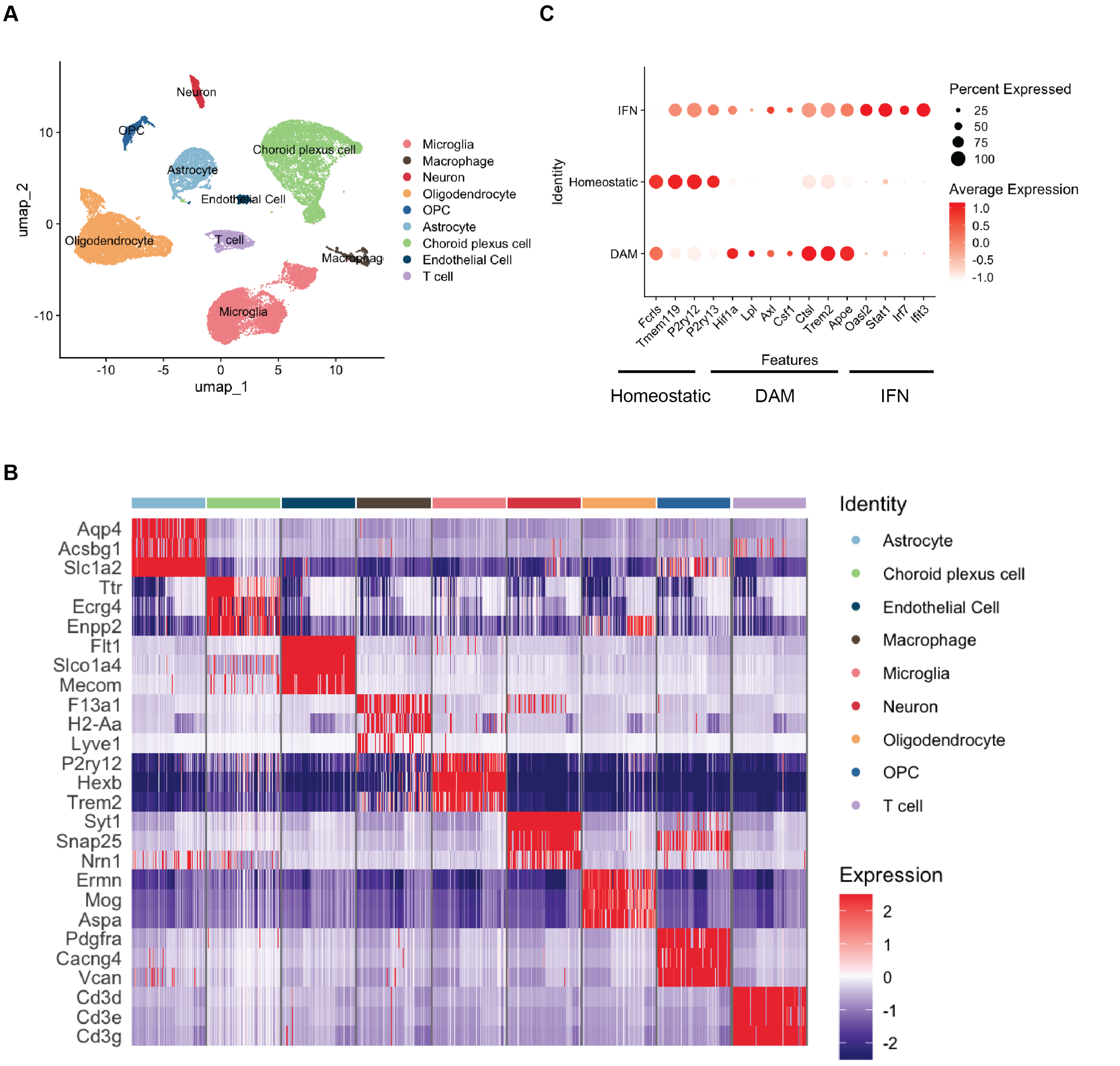


**Supplementary Figure 4 – related to Figure 3**

**(A)** UMAP plots of all single cell and their annotated cell types; **(B)** Gene expression heatmap showing the marker genes for each cluster; **(C)** Dot plot representation of cluster defining genes for each microglial subpopulation.

**Fig. S5.**

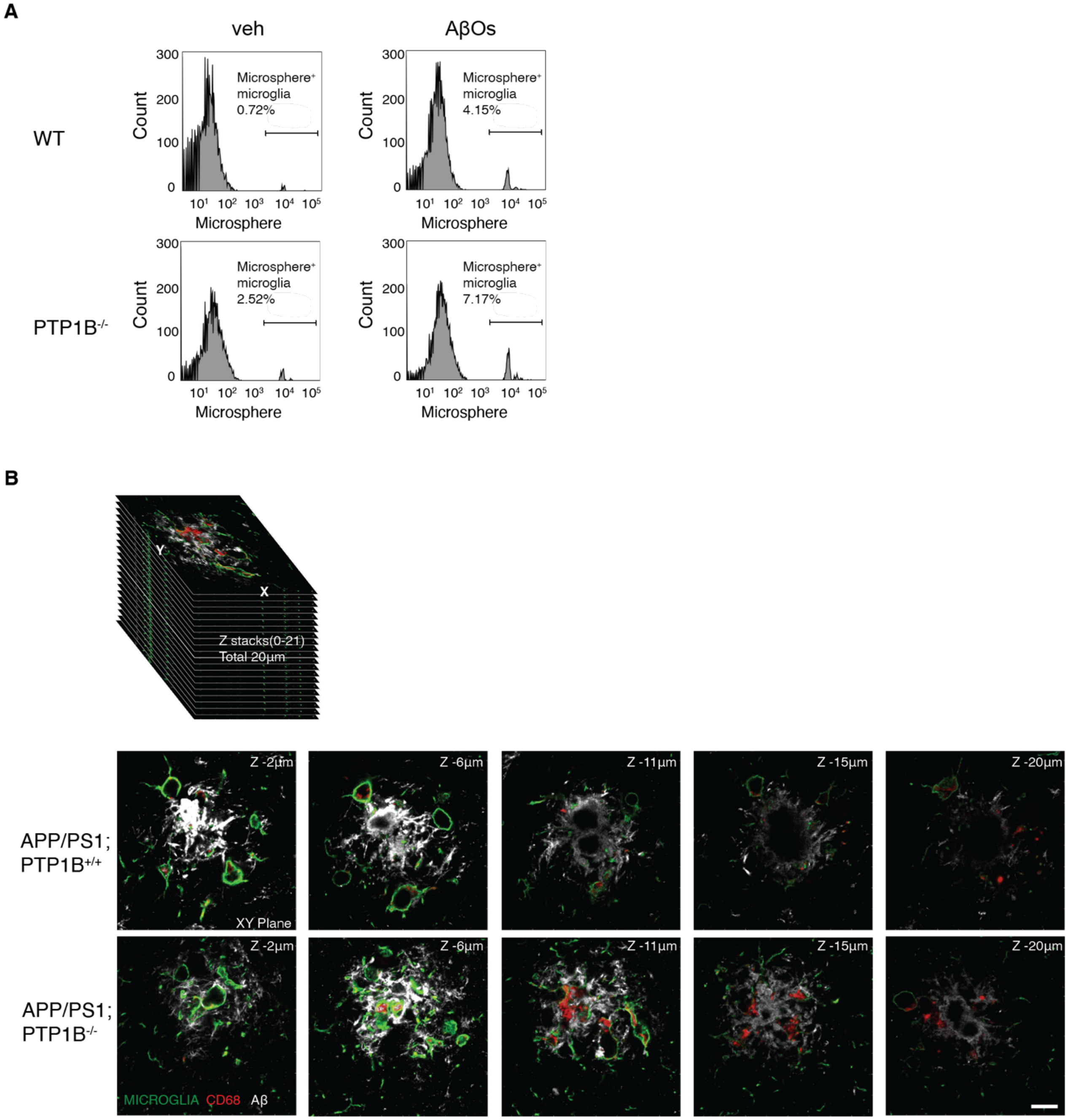


**Supplementary Figure 5 – related to Figure 4**

(**A**) Representative micrographs of WT and PTP1B$^{-/-}$ primary microglia, treated with AβOs, and measurement of phagocytic activity by fluorescence-activated cell sorting (FACS)-based microparticle-uptake assay, quantification was shown in Fig.6C. (**B**)The representative Z-stack pictures for analyzing phagocytosed Aβ by microglia in Figure 4C. XY plane images were serially taken along the z depth (z-stacks: 0-21; total 20 μm) to measure internalized Aβ by phagocytosis. Aβ and microglia, phagocytosis marker are indicated by 6E10, IBA1 and CD68, respectively.

Fig. S6.

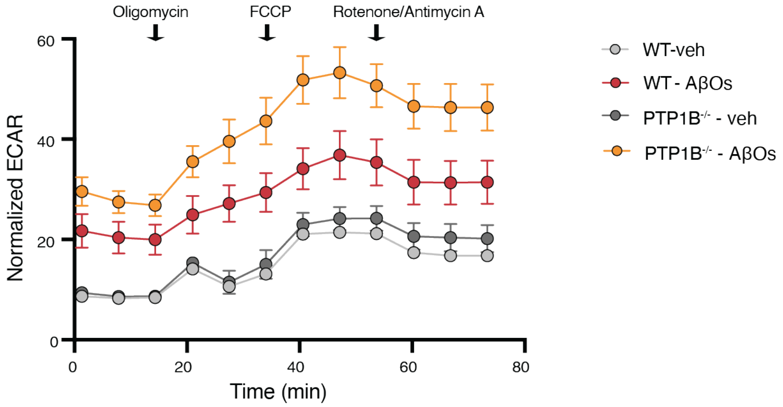


**Supplementary Figure 6 – related to Figure 5**

WT and PTP1B$^{-/-}$ primary microglia were treated with and without AβOs for 24h. After treatment, real time extracellular acidification rate (ECAR) was measured following sequential addition of oligomycin, FCCP, rotenone and antimycin A as illustrated (n = 3 independent experiments, with 5-8 technical replicates per experiment).

**Fig. S7.**

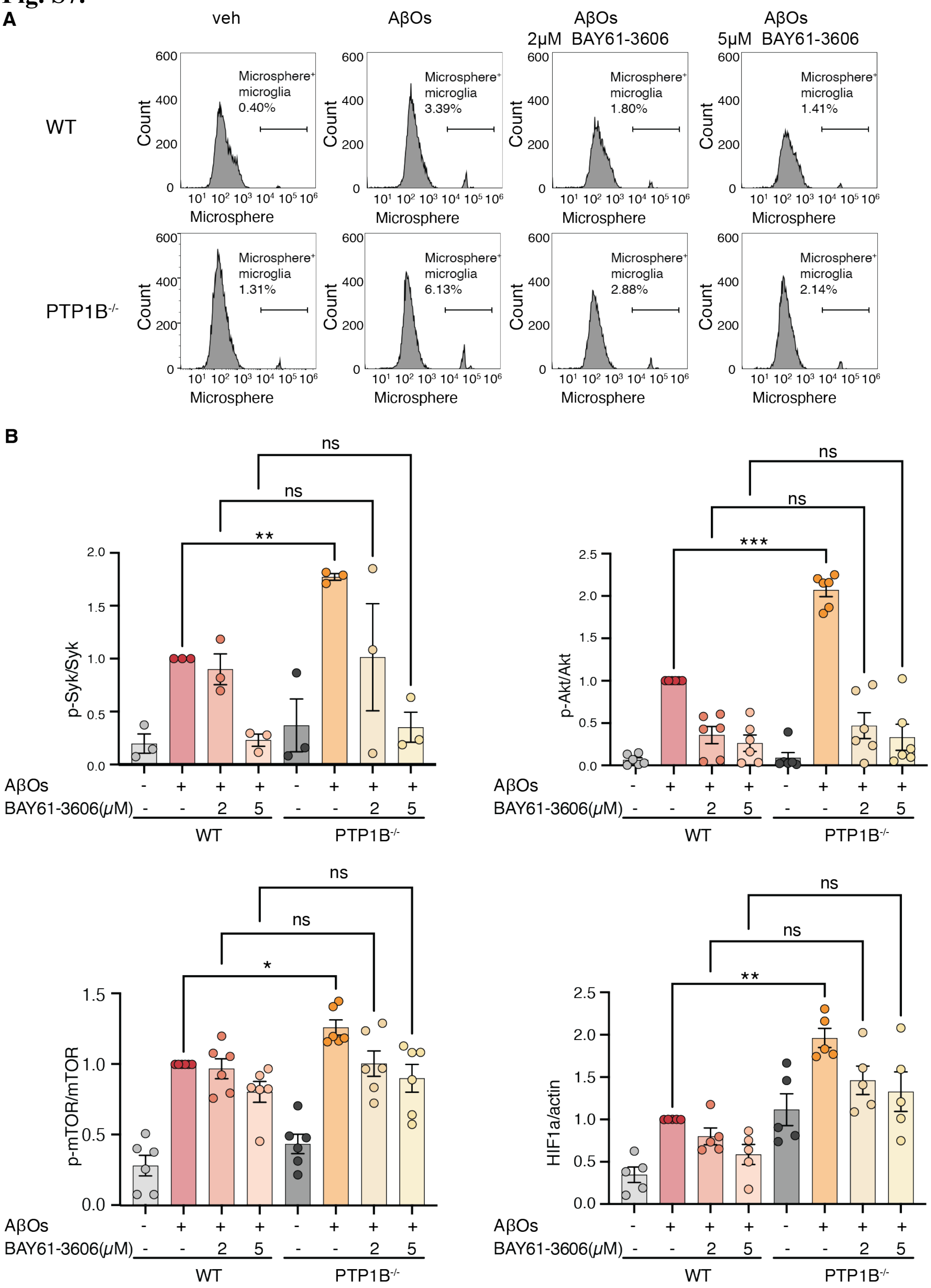


**Supplementary Figure 7 – related to Figure 6**

(**A**) Representative micrographs of WT and PTP1B$^{-/-}$ primary microglia, treated with AβOs or different concentrations of SYK inhibitor BAY-61-3606, and measurement of phagocytic activity by fluorescence-activated cell sorting (FACS)-based microparticle-uptake assay (n = 7 per group), quantification was shown in Fig.6C. (**B**) Quantification of immunoblots shown in Fig.6B, p-SYK, SYK, AKT, p-AKT, mTOR, p-mTOR, HIF1α, and actin in WT or PTP1B$^{-/-}$ microglia treated with AβOs and different concentrations of SYK inhibitor BAY-61-3606.

**Table S1.**

| Antibody name | Catalog no. | RRID | Source |
|---|---|---|---|
| Mouse monoclonal anti-β-Amyloid (clone: 6E10) | 803001 | AB_2564653 | BioLegend |
| Anti-Iba-1 antibody | ab5076 | AB_2224402 | Abcam |
| Anti-CD68 antibody | ab125212 | AB_10975465 | Abcam |
| APC/Cyanine7 anti-mouse/human CD11b Antibody | 101226 | AB_830642 | BioLegend |
| Rabbit monoclonal anti-p-mTOR (Ser2448) | 5536 | AB_10691552 | Cell Signaling Technology |
| Rabbit polyclonal anti-mTOR | 2972 | AB_330978 | Cell Signaling Technology |
| Rabbit polyclonal anti-HIF-1α | NB100-449 | AB_10001045 | Novus |
| Rabbit polyclonal anti-p-AKT (Ser473) | 9271 | AB_329825 | Cell Signaling Technology |
| AKT (pan)(C67E7) Rabbit mAb | 4691 | AB_915783 | Cell Signaling Technology |
| Rabbit monoclonal anti-p-S6 ribosomal protein (Ser235/236) | 4858 | AB_916156 | Cell Signaling Technology |
| Rabbit monoclonal anti-S6 ribosomal protein | 2217 | AB_331355 | Cell Signaling Technology |
| Anti-mouse PTP1B | AF3954 | AB_2174947 | R and D Systems |
| HRP anti-actin | ab20272 | AB_445482 | Abcam |
| SYK (D3Z1E) XP Rabbit Antibody | 13198 | AB_2687924 | Cell Signaling Technology |
| 4G10, monoclonal | 05-321 | AB_309678 | Millipore Sigma |
| Rabbit polyclonal anti-p-SYK (Tyr525/526) | 2711 | AB_2197215 | Cell Signaling Technology |
| HA-Tag | 3724 | AB_1549585 | Cell Signaling Technology |
| FLAG-tag (DYKDDDDK) | 14793 | AB_2572291 | Cell Signaling Technology |
| Anti-PTP1B | 244207 | AB_2877148 | Abcam |
| APP-CTF | A8717 | AB_258409 | Sigma |
| BACE1 | Sc-33711 | AB_626716 | Santa Cruze |
| Presinilin1 | 5643T | AB_10706356 | Cell Signaling Technology |

**Table S1.** Antibodies used in this study
